# VAMP2/Munc18-1 domain 3a interaction controls the nanoscale reorganization underpinning vesicular priming

**DOI:** 10.1101/2023.09.10.556776

**Authors:** Anmin Jiang, Rachel S. Gormal, Ravikiran Kasula, Ailisa Blum, Mercè Salla-Martret, Ye Jin Chai, Tristan P. Wallis, Julie Z. Brouillet, Pranesh Padmanabhan, Saroja Weeratunga, Emma K. Livingstone, Mahdie Mollazade, Merja Joensuu, Ramón Martínez-Mármol, Brett M. Collins, Frédéric A. Meunier

## Abstract

SNARE-mediated secretory vesicle (SV) exocytosis underpins neuronal communication. Munc18-1 orchestrates SNARE complex formation by controlling the opening of syntaxin-1A. How the SV-plasma membrane interface becomes fusion-competent at the nanoscale level is poorly understood. Here, we propose that the interaction of Munc18-1 with VAMP2 during vesicular docking triggers nanoscale re-organization which renders the SV-plasma membrane interface fusion-competent. We identified and mutated key residues in Munc18-1 domain 3a (A297 and T304) hypothesised to impair its interaction with VAMP2. Munc18-1^A297H^, and to a lesser extent Munc18-1^T304H^, constrained SVs on the plasma membrane and reduced stimulated secretion, under re-expression conditions in Munc18-1/2 double knockout neurosecretory cells. Moreover, the de-clustering of Munc18-1 in response to activity was lost for both mutants. The interaction of VAMP2 with the Munc18-1 domain 3a therefore controls the re-organization of the nanoscale environment of the docked SV-plasma membrane interface, fostering syntaxin-1A opening and Munc18-1 release to ensure that SNARE assembly only occurs within the confinement of docked vesicles.

## Introduction

Neurotransmitter release and hormone secretion occur through the process of exocytosis, which involves the fusion of secretory vesicles (SVs) with the plasma membrane in neurons and neurosecretory cells. How the plasma membrane becomes conducive to fusion of SVs is unknown. However, we know that stimulation elicits a discrete number of molecular steps that enables the establishment of multiple fusion pores on the plasma membrane. Understanding how molecules specialized in this process can assemble a fusion-competent interface between the plasma membrane and SVs at the nanoscale is therefore critical. SVs undergo stepwise docking and priming to acquire the ability to fuse with the plasma membrane in a Ca^2+^-dependent manner ^1^. The ultimate fusion step is mediated by protein-protein interactions between soluble N-ethylmaleimide-sensitive factor attachment protein receptors (SNAREs) and the Sec/Munc18 family protein Munc18-1. The SNAREs can be subdivided into categories including the target membrane-associated t-(or Q-) SNAREs (syntaxin-1A, SNAP25), and the vesicle-associated v-(or R-) SNARE (vesicle-associated membrane protein 2 (VAMP2) or synaptobrevin) ^1^. The formation of the SNARE complex is essential to overcome the energy barrier associated with the fusion of two negatively charged phospholipid bilayers, such as the plasma membrane and the SV membrane ^2^. This process is mediated by the zippering of the t- and v-SNAREs into a four-stranded parallel coiled-coil ^3,4^. Several regulatory proteins are necessary to control SNARE-mediated membrane fusion. Munc18-1 is one such protein that binds directly to syntaxin-1A, holding it in a closed conformation in the absence of other factors, which inhibits syntaxin-1A from interacting with the other SNARE proteins thus preventing uncontrolled SV fusion ^5–10^. In addition to this, Munc18-1 also plays a critical role in the docking and priming of SVs ^2,11–13^. Further, Munc18-1, together with Munc13-1, orchestrates the opening of syntaxin-1A to allow efficient SNARE complex assembly ^14–16^.

The role of various Munc18-1 domains in regulating SNARE assembly has been debated. Biochemical experiments have revealed that an interaction occurs between the Munc18-1 domain 1 hydrophobic pocket and the N-terminal region of syntaxin-1A ^8,17–19^. Although this interaction was deemed to play a role in SNARE complex formation during the priming of vesicles, a mutation which blocks this interaction, Munc18-1^F115E^, did not result in any significant defects in exocytosis in neurosecretory cells (Malintan et al., 2009), and in neurons ^20,21^. Other reported mutations in the Munc18-1 domain 1 all result in a defect in syntaxin-1A transport to the plasma membrane ^22^. The Munc18-1 domain 3a can transition from a closed to an extended conformation, with the latter being incompatible with binding to the closed conformation of syntaxin-1A ^23,24^.This suggests that a bent-to-extended conformational change in the Munc18-1 domain 3a could trigger the opening of syntaxin-1A at the plasma membrane to induce vesicle priming ^23,24^. Indeed, a Munc18-1 deletion mutant lacking 17 residues of the domain 3a loop engineered to block the conformational loop change (Munc18-1^Δ317–333^), prevented the engagement of syntaxin-1A into the SNARE complex, but did not affect its transport to the plasma membrane ^25,26^. The activity-dependent release of Munc18-1 from the confinement of nanoclusters at the inner leaflet of the plasma membrane was also lost for the Munc18-1^Δ317–333^ mutant ^25^. Previous findings suggest that Munc18-1 binds to other SNARE proteins in the presence of open syntaxin-1A ^17,27^ and that the Munc18-1 domain 3a could act as a template for SNARE proteins to initiate the four helix coiled-coil zippering ^28^. This concept was supported by a study whereby Vps33, a Munc18-like protein in yeast, was suggested to act as a template for Vam3 (the yeast homolog of syntaxin-1A) and Nyv1 (yeast VAMP2-like protein) to generate partially zippered SNARE assembly intermediates ^29^. These findings suggest the existence of an intermediary SNARE complex via the interaction of VAMP2 and Munc18-1 during SV fusion. It has been proposed that open syntaxin-1A molecules form bunch-like clusters on the plasma membrane ^30–32^. However, considering that a constitutively open syntaxin-1B mutant causes uncontrolled SV fusion and seizures ^7^, we hypothesize that syntaxin-1A only opens in the context of vesicular docking upon transient binding of VAMP2 to the Munc18-1 domain 3a hinge-loop. This binding could stabilize the extended conformation of the Munc18-1 domain 3a loop, thereby facilitating the opening of syntaxin-1A and productive SNARE complex formation.

In the current study, we analyzed the X-ray crystal structure and sequence alignments of the yeast Vps33/Nyv1 complex ^29^ as well as the recently published cyro-EM structure of *Rattus norvegicus* of the VAMP2/Munc18/Sx1a complex ^33^ to identify potential residues that form the interface between Munc18-1 and VAMP2. From this, we generated two VAMP2 binding-deficient Munc18-1 mutants (A297H and T304H), and studied their effects in neurosecretory PC12 cells which were engineered to knock out Munc18-1 and -2 using the CRISPR-Cas9 system ^34^. In Munc18-1 and -2 double-knockout PC12 (DKO-PC12) cells, SVs lose their ability to undergo docking and fusion. These effects were rescued by the re-expression of Munc18-1^WT^. However, re-expression of Munc18-1^A297H^ led to a ‘super-docking’ phenotype, whereby vesicles exhibited significantly reduced mobility at the plasma membrane at rest, and no further immobilization was observed in response to secretagogue stimulation. In addition, the previously observed activity-dependent exit of Munc18-1^WT^ molecules from plasma membrane nanodomains ^25^ was not detected in either VAMP2 binding-deficient mutants. The inability of Munc18-1 mutant A297H to rescue the activity-dependent membrane re-organization, was further observed by tracking syntaxin-1A molecules ^25^. These results suggest that both Munc18-1^A297H^ and (to a lesser extent) Munc18-1^T304H^ arrest SVs in a non-functional docked state, and that VAMP2 binding to Munc18-1 triggers the exit of Munc18-1 from nanodomains, as well as the opening of syntaxin-1A, which enables efficient SNARE complex assembly.

## Results

### Munc18-1 domain 3a residues A297 and T304 mediate VAMP2 binding in neurosecretory cells

Analysis of the Munc18/Vamp2/Sx1a complex structure determined by cryoEM ^33^ and comparison with the yeast Vps33/Nyv1 complex structure^29^ identified two potential Munc18-1 residues (T304 and A297) that form the interface between Munc18-1 and R-SNARE VAMP2 and may be generally involved in assembly with the bound Q-SNARE Sx1a. We hypothesized that mutating the T304 and A297 residues to bulky histidines would disrupt VAMP2 binding via steric hindrance (Fig. 1A). To determine the effect of these Munc18-1 A297H and T304H mutations on the interaction of Munc18-1 with VAMP2, we performed pulldown assays with recombinantly expressed and purified Munc18-1^WT^-His, Munc18-1^A297H^-His or Munc18-1^T304H^-His and VAMP2-GST (glutathione S-transferase). We could detect binding of VAMP2 to Munc18-1^WT^ as previously found ^28^, with either 1 h incubation at pH 6.5 or overnight incubation. However, the signal was faint suggesting that their interaction *in vitro* is weak or transient (Fig. 1B-C and S1). Although the binding of Munc18-1^T304H^ to VAMP2 was similar to that of Munc18-1^WT^, we could hardly detect any binding between VAMP2 and the Munc18-1^A297H^ mutant even under the prolonged incubation conditions (Fig. 1B-C).

**Figure 1:**
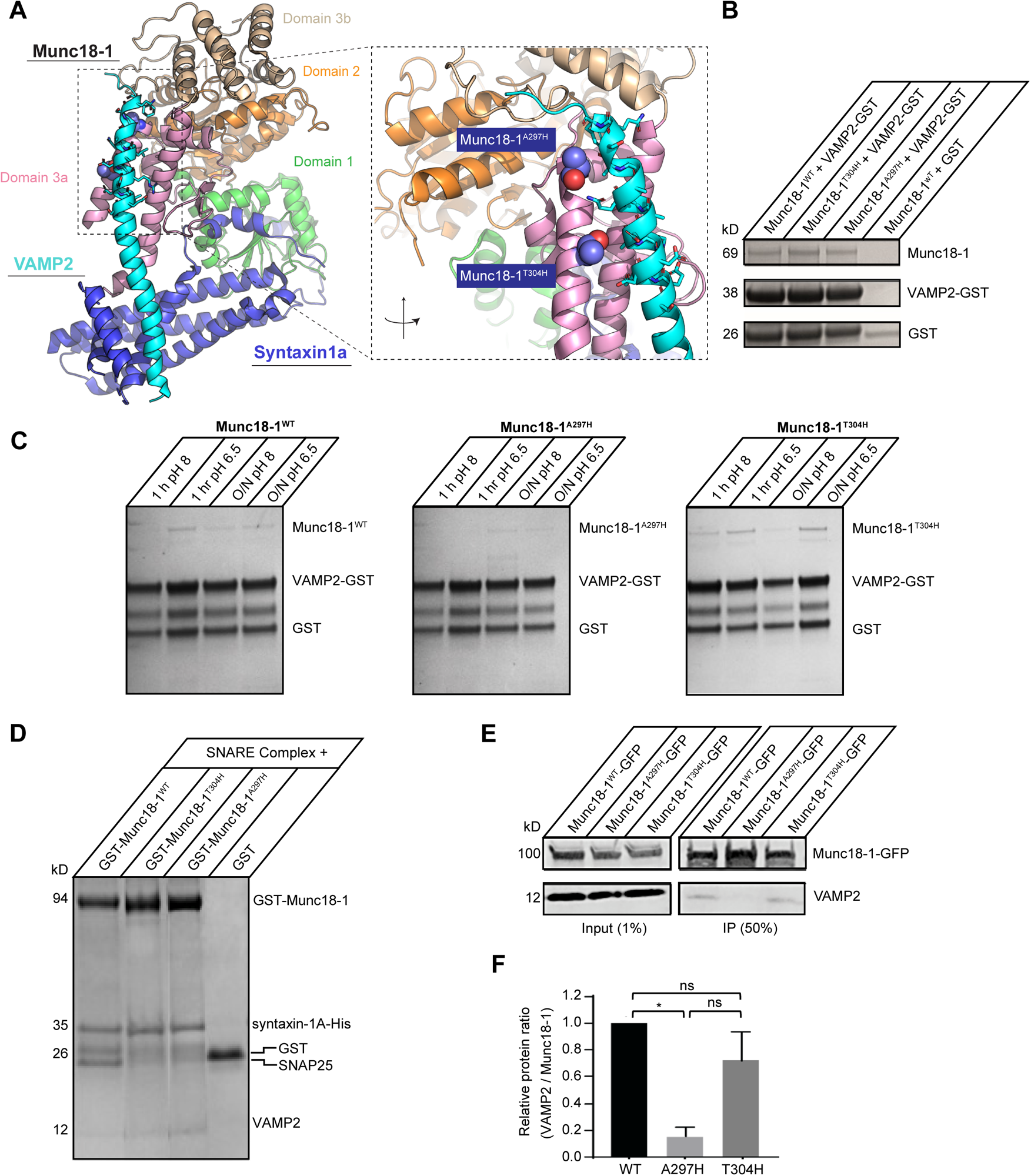
Munc18-1 domain 3a residues A297 and T304 control its interaction with VAMP2 in neurosecretory cells. **(A)** CryoEM structure of *Rattus norvegicus* VAMP2/Munc18-1/Syntaxin1 (PDB ID 7UDC) ^33^ illustrates Munc18-1 domain 3b (brown) and domain 3a (pink) forming a template for VAMP2 (cyan). Based on this model, amino acid residues Ala297 and Thr304 in Munc18-1, were predicted to be potential binding sites for VAMP2. VAMP2 binding-deficient Munc18-1 mutants (Munc18-1^A297H^ and Munc18-1^T304H^) were generated accordingly. **(B)** Recombinantly expressed Munc18-1^WT^-His, Munc18-1^T304H^-His or Munc18-1^A297H^-His was incubated with VAMP2-GST and glutathione agarose beads overnight at pH 8.0, immobilized on glutathione agarose resin, analyzed by SDS-PAGE and the gels stained with Coomassie blue. As a control, Munc18-1^WT^-His was also incubated with GST alone. The results were reproduced in three independent experiments. **(C)** The experiment shown was repeated for Munc18-1^WT^-His, Munc18-1^T304H^-His and Munc18-1^A297H^-His using different incubation times (either 1 h or overnight) and pH values (either 6.5 or 8.0). **(D)** Preformed SNARE complexes (syntaxin-1A-His, SNAP25, VAMP2) were incubated with either GST-Munc18-1^WT^, GST-Munc18-1^T304H^, GST-Munc18-1^A297H^ or GST alone (control), and glutathione sepharose beads for 30 min. Bound proteins were analyzed by SDS-PAGE and the gels stained with Coomassie blue. The results were reproduced in two independent experiments. **(E)** Munc18-1^WT^-emGFP, Munc18-1^A297H^-emGFP or Munc18-1^T304H^-emGFP was expressed in DKO-PC12 cells and immunoprecipitated with GFP-Trap beads. **(F)** Relative protein ratios of VAMP2 / Munc18-1-emGFP were quantified from the co-immunoprecipitations in (E) from three independent experiments. Results in (F) are expressed as mean ± SEM. A Kruskal Wallis test with multiple comparisons was performed (ns, not significant; * p < 0.05).

We then tested whether these mutations affected the interaction between Munc18-1 and the preformed SNARE complexes (syntaxin-1A-His + VAMP2 + SNAP25) ^8,20,35^. We detected binding of Munc18-1 to preformed SNARE complexes after 30 min incubation. Immunoprecipitation revealed that GST-tagged Munc18-1^WT^, Munc18-1^A297H^ and Munc18-1^T304H^ were all able to bind to the preformed SNARE complexes (Fig. 1D).

As the *in vitro* experiments indicated that the interaction of Munc18-1 and VAMP2 is weak and might require additional proteins, we next investigated the effects of the Munc18-1 A297H and T304H mutations on the interaction with VAMP2 in neurosecretory cells. To this end, we generated Munc18-1 and -2 double knockout (DKO)-PC12 cells using the CRISPR-Cas9 system (Fig. S2). Previously validated Munc18-1 knockout cells (Chai et al. 2014 *JCB*) were used as a template to generate Munc18-1/2 DKO-PC12 cells. The frameshift induced by CRISPR-Cas9 resulted in an early stop codon in Munc18-2 at the level of exon 3 (Fig. S2) as verified by Sanger sequencing of mRNA following RT-PCR. Similarly to the previously published Munc18-1/2 knockdown cells ^25,26^, the fusion of SVs was severely impaired in the DKO-PC12 cells, which was rescued by the re-expression of Munc18-1^WT^-mEos2 (Fig. S2). Further, we examined whether Munc18-1^WT^-emGFP, Munc18-1^A297H^-emGFP or Munc18-1^T304H^-emGFP affected the expression and targeting of syntaxin-1A to the plasma membrane in DKO-PC12 cells. In these DKO-PC12 cells, like in DKD-PC12 cells ^26,36^. Syntaxin-1A transport to the plasma membrane was severely impaired and rescued upon re-expression of Munc18-1^WT^. We found that all constructs were capable of rescuing syntaxin-1A localization on the plasma membrane (Fig. S3) indicating that these residues do not play a role in Syntaxin-1A targeting to the plasma membrane.

To further assess the binding of Munc18-1 to VAMP2, the DKO-PC12 cells were transfected with Munc18-1^WT^-emGFP, Munc18-1^A297H^-emGFP or Munc18-1^T304H^-emGFP which were then immunoprecipitated using GFP-trap. VAMP2 was co-immunoprecipitated with immobilized Munc18-1^WT^-emGFP. However, interaction with VAMP2 was dramatically reduced for the Munc18-1^A297H^-emGFP mutant (Fig. 1 D, E). In contrast to this, the level of interaction of Munc18-1^T304H^-emGFP with VAMP2 was only slightly lower than that of Munc18-1^WT^, suggesting that this mutation has a much lower impact on the transient VAMP2/Munc18-1 interaction.

Although the Munc18-1 A297H and T304H mutations did not have a major impact on the interaction of purified Munc18-1 with VAMP2 *in vitro* (Fig. 1), the A297 residue was critical for the interaction between the two proteins in neurosecretory cells, with the T304 being less important. Our results suggest that the binding of VAMP2 to Munc18-1 likely involves additional proteins that are only present in the cellular context. This is in good agreement with a recent report also suggesting the involvement of Munc13 in this process^37^.

### VAMP2 binding-deficient Munc18-1 mutants exhibit a super-docking phenotype

Munc18-1 controls the docking of SVs in neurosecretory cells ^12,22^ and compromising domain 3a hinge-loop function has been shown to immobilize SVs on the plasma membrane resulting in a ‘super-docking’ phenotype ^25^. We tested whether the VAMP2 binding-deficient Munc18-1 mutants also affected the mobility of SVs by total internal reflection fluorescence (TIRF) microscopy. Neuropeptide Y (NPY) is packaged in large dense core vesicles in PC12 cells and is therefore used as a SV marker ^38^. Taking advantage of this, DKO-PC12 cells were co-transfected with NPY-emGFP and either Munc18-1^WT^-mCherry, Munc18-1^A297H^-mCherry, Munc18-1^T304H^-mCherry or pCMV empty vector to determine the mobility of SVs on the plasma membrane in response to stimulation (2 mM BaCl_2_) (Fig. 2 A-C). As expected, in cells expressing Munc18-1^WT^-mCherry, the NPY-emGFP positive SVs became more confined in response to stimulation compared to cells which did not express any Munc18-1 (pCMV) (Fig. 2 D). This is in good agreement with a role of Munc18-1^WT^ in SV docking ^12,25,26,39^. In sharp contrast, we could not detect this activity-dependent immobilization of SVs in cells expressing Munc18-1^A297H^-mCherry or Munc18-1^T304H^-mCherry. Surprisingly, SVs in these cells were already much more immobile at rest and their mobility did not decrease further in response to secretagogue stimulation (Fig. 2 D). We therefore concluded that Munc18-1^A297H^-mCherry and Munc18-1^T304H^-mCherry promote a super-docking phenotype in which vesicles are already docked prior to stimulation and remain unchanged following stimulation. This effect was more pronounced in cells expressing Munc18-1^A297H^-mCherry than in cells expressing Munc18-1^T304H^-mCherry (Fig. 2 D) suggesting that the latter affected docking to a lesser extent and consistent with the more pronounced VAMP2-binding effect of the A297H mutant.

**Figure 2:**
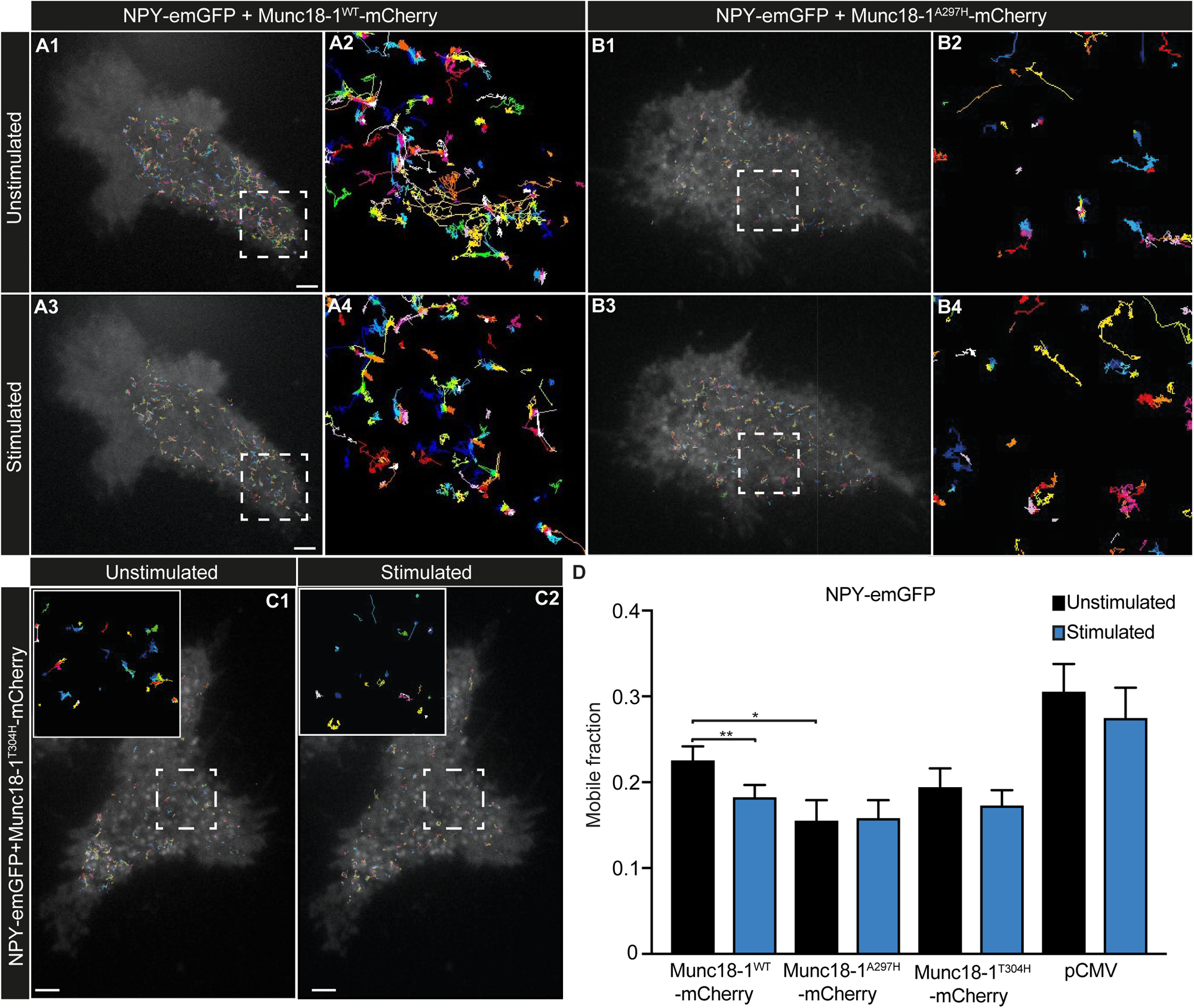
Mutation of Munc18-1 domain 3a residues A297 and T304 causes super-docking of secretory vesicles. **(A-D)** DKO-PC12 cells were co-transfected with NPY-emGFP (SV marker) and either (**A**) Munc18-1^WT^-mCherry, **(B)** Munc18-1^A297H^-mCherry or **(C)** Munc18-1^T304H^-mCherry, and imaged by TIRF microscopy at 20 Hz before and after secretagogue stimulation using BaCl_2_ (2 mM). pCMV empty vector was used as a control. The trajectories of single NPY-emGFP positive SVs for each condition are superimposed on the corresponding Munc18-1-mCherry TIRF image showing the level of Munc18-1-mCherry expression. (Insert scale bars: 2 µm). The respective single NPY-emGFP SV trajectories from the enlarged inserts are shown before and after stimulation in **(A2, 4, B2, 4 and C1, 2)** (insert scale bars: 500 nm). **(D)** Quantification of the mobile fractions of NPY-emGFP SVs before and after stimulation, calculated from the distribution of diffusion histograms using PALM tracer. Results in (D)are expressed as mean ± SEM. A paired student’s *t*-test (before and after stimulation) and unpaired student’s *t*-test to compare the mobile fraction of cells expressing the various Munc18-1 constructs or pCMV were performed (* p < 0.05; ** p < 0.01; *** p < 0.001).

### NPY release is decreased in VAMP2 binding-deficient Munc18-1 mutants

We and others have previously demonstrated that the Munc18-1 domain 3a hinge-loop is essential to rescue SV fusion in the absence of Munc18-1/2 ^24–26,40^. Therefore, we next investigated the effect of the VAMP2 binding-deficient Munc18-1 mutants on the fusion of SVs. DKO-PC12 cells were co-transfected with VAMP2-pHluorin and either Munc18-1^WT^-mCherry, Munc18-1^A297H^-mCherry or Munc18-1^T304H^-mCherry and were stimulated with 2 mM BaCl_2_ and visualized using TIRF microscopy. The pH-sensitive GFP variant pHluorin is quenched in the acidic environment of SVs and becomes unquenched upon fusion of the SVs with the plasma membrane ^41^. This unquenching leads to an abrupt increase in fluorescence, which allows the quantification of SV exocytosis. A representative SV fusion event is shown in Fig. 3 A-B. The DKO-PC12 cells transfected with Munc18-1^WT^-mCherry showed a drastic increase in VAMP2-pHluorin fluorescence intensity following stimulation (Fig. 3 A-B and E-F). This represents the result of all the fusion events leading to VAMP2-pHluorin translocation and unquenching at the plasma membrane. This overall plasma membrane fluorescence increase was significantly dampened in DKO-PC12 cells expressing Munc18-1^A297H^-mCherry (Fig. 3 C, E-F). It was also reduced in Munc18-1^T304H^-mCherry expressing DKO-PC12 cells, although the trend did not reach significance (Fig. 3 D-F). Similar results were obtained using depolarization (high K^+^) buffer to stimulate DKO-PC12 cells (Fig. S4). These results indicate that Munc18-1 residue A297, and to a lesser extent residue T304, are critical for SV release. This is in accordance with our co-immunoprecipitation results, which showed that the VAMP2 interaction was reduced for Munc18-1^A297H^ but less so for Munc18-1^T304H^ so in the mutant (Fig. 1 E-F). Similarly, another Munc18-1^L348R^ mutant which does not interact with VAMP2 has also been reported to cause decreased vesicle fusion ^24,28^.

**Figure 3:**
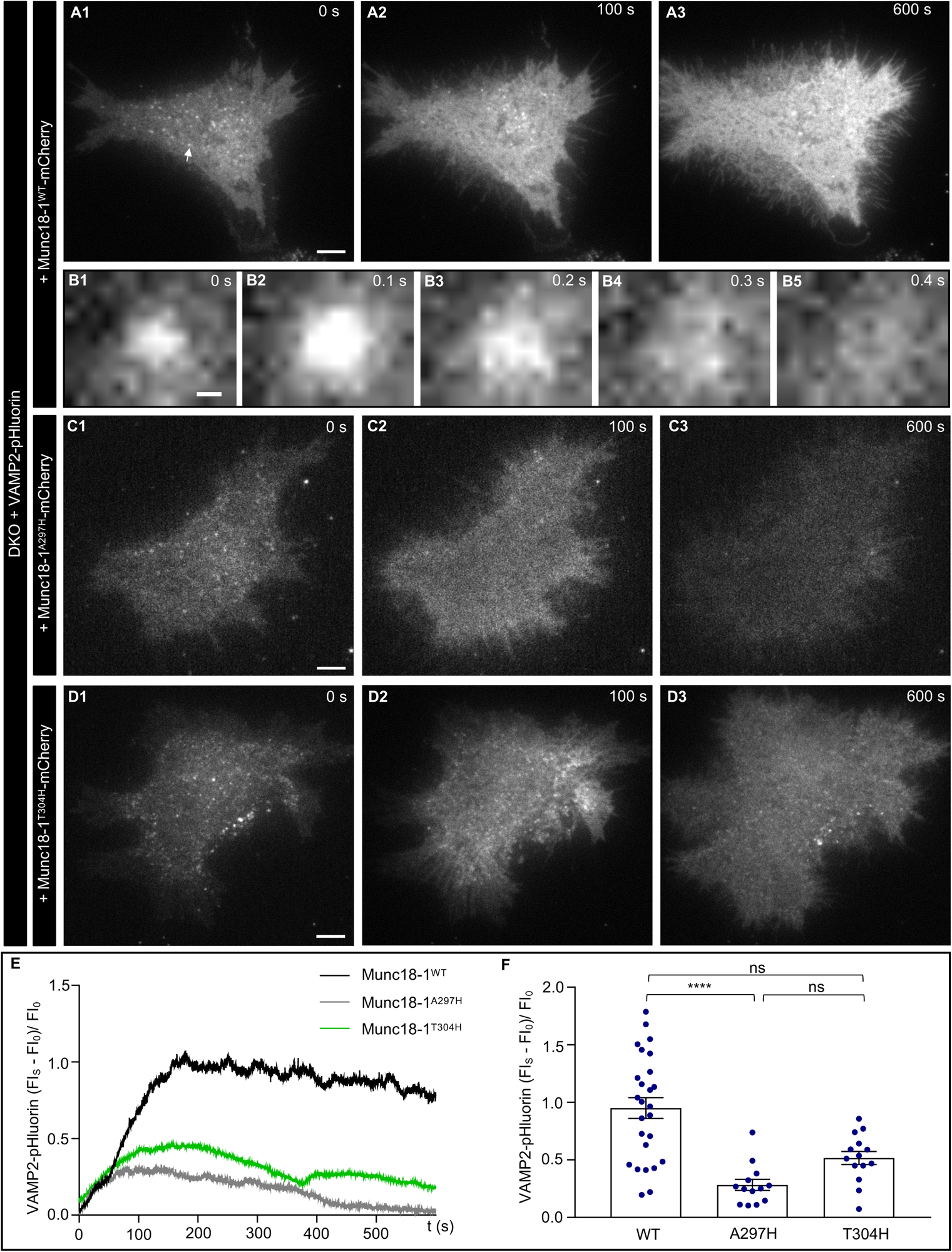
Secretory vesicle exocytosis is decreased in VAMP2 binding-deficient Munc18-1 mutants. **(A-D)** DKO-PC12 cells were co-transfected with VAMP2-pHluorin and either **(A,B)** Munc18-1^WT^-mCherry, **(C)** Munc18-1^A297H^-mCherry or **(D)** Munc18-1^T304H^-mCherry (scale bars: 5 µm). SV release was determined by the fluorescence intensity (FI) increase of VAMP2-pHluorin after stimulation. Cells were imaged at 10 Hz before stimulation and immediately after stimulation (2 mM BaCl_2_) to elicit secretion. The arrow in (A1) indicates a VAMP2-pHluorin-positive SV, which is enlarged in (B) as a timelapse series showing an SV fusing with the plasma membrane. (E) Traces of the VAMP2-pHluorin FI fold change after stimulation (FI_S_) normalized by the FI before stimulation (FI_0_) from one representative cell (FI_S_ - FI_0_)/FI_0_). (**F**) The FI fold change at 200 s after stimulation. Results are expressed as mean ± SEM. A Kruskal-Wallis test with multiple comparisons was performed (ns, not significant; **** p < 0.0001).

### The interaction of Munc18-1 with VAMP2 controls the activity-dependent release of Munc18-1 from nanoclusters on the plasma membrane

Munc18-1 is organized in nanoclusters on the plasma membrane, which overlap with SNAP25 and syntaxin-1A nanoclusters ^42,43^. We have previously shown that following stimulation, Munc18-1 must be released from these nanoclusters to ensure efficient SV exocytosis ^25^. To analyze the influence of VAMP2 binding on the nanoscale organization of Munc18-1, we performed spatiotemporal cluster analysis and assessed the dynamic re-organisation of Munc18-1 in response to stimulation. Specifically, we used an adapted spatiotemporal DBSCAN (density-based spatial clustering of applications with noise) approach called BOOSH, which enabled us to analyse the dynamics of cluster size and density in response to stimulation in cells expressing VAMP2-bindingdeficient Munc18 mutants.

BOOSH is an experimental derivative of a spatiotemporal clustering algorithm ^44^ which uses a modified 3D DBSCAN ^45^ approach to establish whether trajectories are clustered in both space and time. In DBSCAN, points are considered clustered if a threshold number of points (MinPts) are within a given spatial proximity radius (ε). BOOSH extends this into the time dimension by establishing a time window (w) within which two trajectories would be considered concurrent. Each detection point is converted from [x,y,t]→[x,y,z] where z = t(ε/w). A trajectory is considered as potentially clustered if any of its individual detections lie within ε of MinPts points from at least two other trajectories. Spatiotemporal clustering metrics are subsequently recovered using a GUI framework (Fig. 4 A).

**Figure 4:**
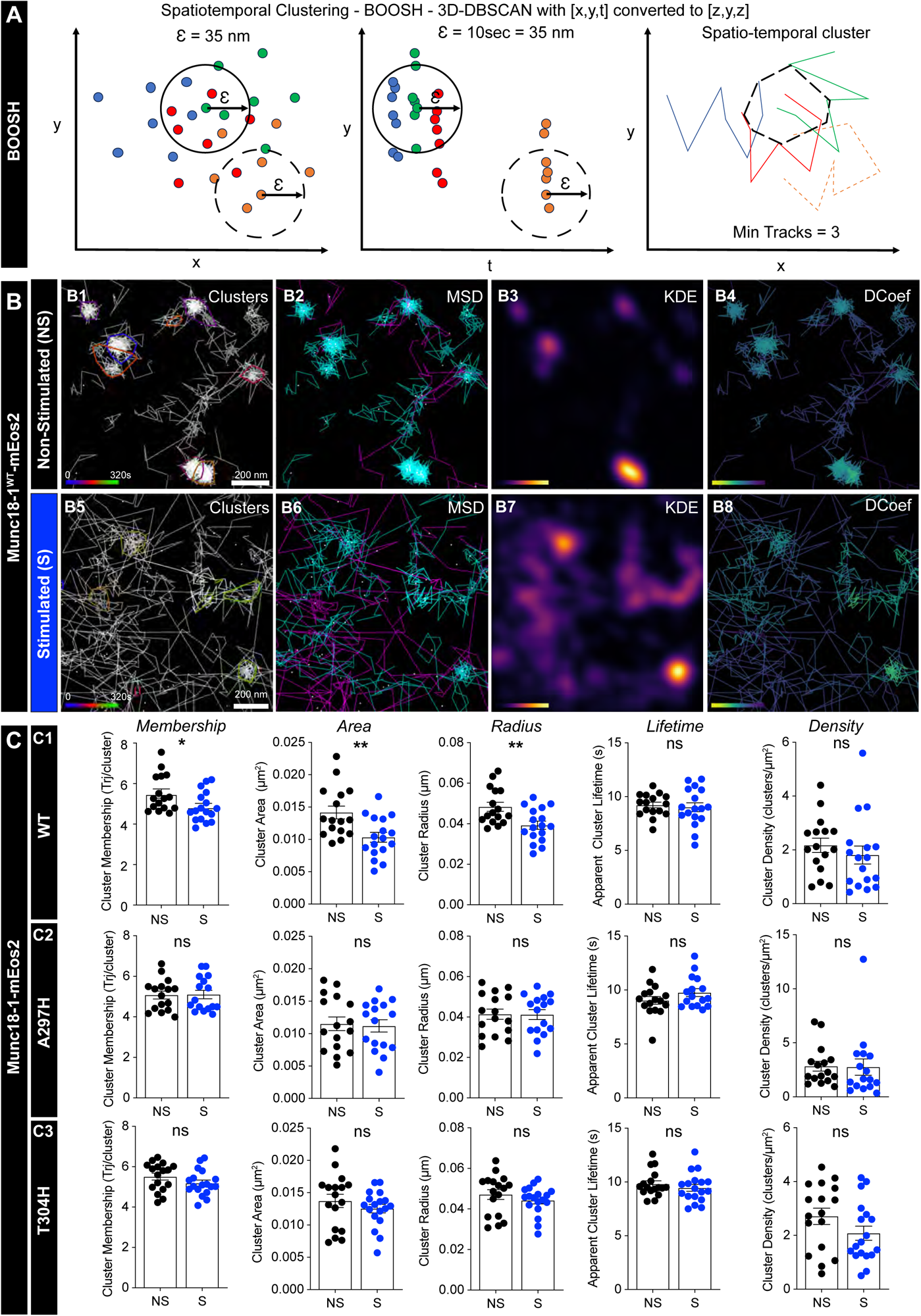
Munc18-1 binding to VAMP2 controls the activity-dependent release of Munc18-1 from nanoclusters on the plasma membrane, revealed using spatiotemporal cluster analysis. **(A)** Schematic diagram of the BOOSH analysis principle. **(B-C)** DKO-PC12 cells expressing **(B, C1)** Munc18-1^WT^-mEos2, **(C2)** Munc18-1^A297H^-mEos2 or **(C3)** Munc18-1^T304H^-mEos2 were imaged at 50 Hz in either non-stimulated (NS) or stimulated (S) conditions (2 mM BaCl_2_). BOOSH analysis was performed on single particle trajectories to determine the effect of stimulation on cluster size and dynamics. Representative Munc18-1^WT^-mEos2 **(B1, B5)** cluster trajectory, **(B2, B6)** MSD, **(B3, B7)** Kernal density estimation and **(B4, B8)** diffusion coefficient maps in non-stimulated and stimulated conditions respectively. Cluster Membership (trajectories/cluster), Cluster Area (um^2^), Cluster Radius (μm), Apparent Cluster lifetime (s) and Cluster Density (clusters/um^2^) were assessed for each condition. Results in (C) are expressed as mean ± SEM, with n = 16-18 cells for each condition. An unpaired student’s *t*-test was performed (ns, not significant; * p < 0.05; ** p < 0.01).

We performed single particle tracking photoactivated localization microscopy (sptPALM) experiments in live DKO-PC12 neurosecretory cells following re-expression with either Munc18-1^WT^, Munc18-1^A297H^ or Munc18-1^T304H^ genetically tagged with the photoconvertible mEos2, in the presence or absence of stimulus (2 mM BaCl_2_). Using spatiotemporal cluster analysis, we found that Munc18-1 forms clusters in unstimulated Munc18-1^WT^-expressing DKO-PC12 cells, with a radius of approximatively 40-50 nm (Fig. 4 B, C1). Upon stimulation, the radius, area and membership (number of trajectories per cluster) were significantly reduced (Fig. 4 B, C1). This decrease in Munc18-1 cluster size and membership is consistent with previous reports ^25^. In contrast, there was no change in clustering metrics for either of the two VAMP2 binding-deficient mutants in response to stimulation (Fig. 4 C2-3). However, we noted that at rest, the values for cluster radius and area were lower for the A297H mutant compared to the T304H mutant and WT Munc18-1 (Fig. 4 C), suggesting that Munc18-1^A297H^ has a stronger clustering phenotype. We observed that Munc18-1 clusters had an apparent lifetime of 8-10s regardless of mutations or stimulation (Fig. 4 C) suggesting that duration of the lateral trap underpinning Munc18-1 clustering is likely independent of its interaction with VAMP2 and the mechanism of fusion.

To further examine this clustering phenotype, we performed fixed PALM and DBSCAN analysis. While the cluster size and total detections of Munc18-1^WT^ on the membrane was also reduced following stimulation, this trend was not statistically significant (Fig. S5). Overall, our fixed and live cluster analysis confirms that Munc18-1^WT^ is released from clusters following stimulation and that mutations affecting VAMP2 interaction prevents this exit.

### The activity-dependent increase in Munc18-1 mobility is dependent on VAMP2 interaction

It has been proposed that Munc18-1 could act as a template for initiating a partial zippering of syntaxin-1A and VAMP2 ^29^. To analyze the effect of VAMP2 binding on the mobility of Munc18-1 molecules, we performed single molecule mobility analysis of the sptPALM data. Conventional TIRF microscopy showed that Munc18-1^WT^-mEos2 is distributed non-uniformly across the plasma membrane (Fig. 5 A1), the sptPALM super-resolved intensity, trajectory and diffusion coefficient maps further highlighted the clustered distribution of Munc18-1^WT^-mEos2 on the plasma membrane (Fig. 5 A2-A4). Individual Munc18-1^WT^-mEos2, Munc18-1^A297H^-mEos2 and Munc18-1^T304H^-mEos2 molecules were tracked to analyse their nanoscale mobility, including mean square displacement (MSD) (Fig. 5 B, E, H) and frequency distribution of diffusion coefficient (D) (expressed as Log_10_D) (Fig. 5 C, F, I) and derived mobile to immobile ratio (Fig. 5 D, G, J). Our analysis revealed two distinct populations of Munc18-1^WT^-mEos2 molecules, a mobile population and an immobile population (Fig. 5 A2-4, C, D).

**Figure 5:**
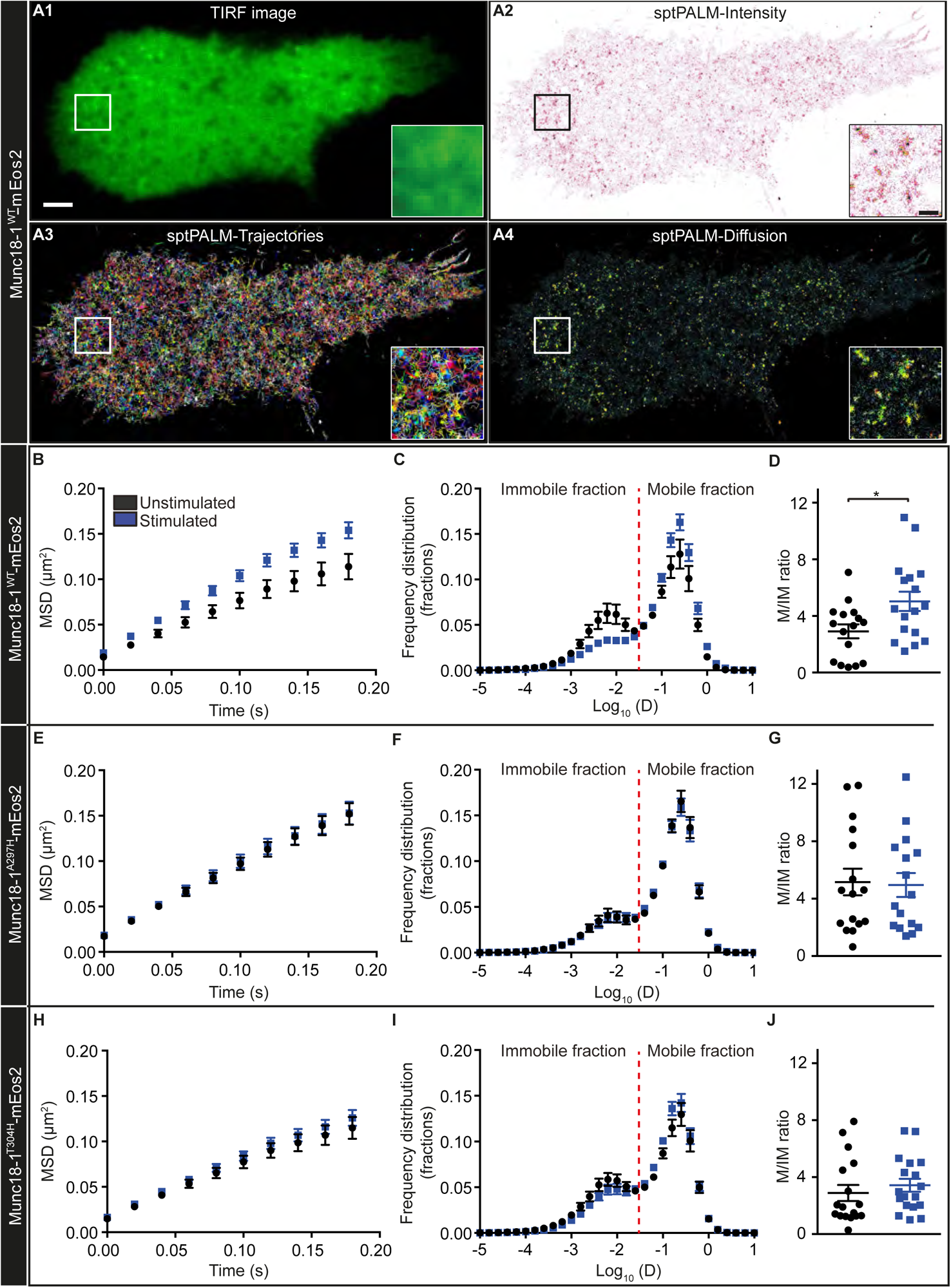
Munc18-1 binding to VAMP2 controls the activity-dependent mobility increase of Munc18-1. DKO-PC12 cells expressing either Munc18-1^WT^-mEos2, Munc18-1^A297H^-mEos2 or Munc18-1^T304H^-mEos2 were imaged by TIRF microscopy at 50 Hz in unstimulated or stimulated conditions (2 mM BaCl_2_). Representative **(A1)** TIRF image (scale bar: 2 µm) and corresponding **(A2)** sptPALM average intensity map, **(A3)** trajectory map (trajectory color-coding in arbitrary units) and **(A4)** diffusion coefficient map, of a DKO-PC12 cell expressing Munc18-1^WT^-mEos2 (detections ranged from 10^-5^ µm^2^/s to 10^1^ µm^2^/s). The enlarged inserts from the region of interest are shown in the respective panels (scale bar: 500 nm). Quantification of single **(B-D)** Munc18-1^WT^-mEos, **(E-G)** Munc18-1^A297H^-mEos2 and **(H-J)** trajectories shown as **(B, E, H)** the average mean square displacement (MSD) as a function of time, **(C, F, I)** the frequency distribution of the diffusion coefficient ^60^ and **(D, G, J)** the mobile/immobile (M/IM) ratio in both unstimulated and stimulated conditions. Results in (D, G and J) are expressed as mean ± SEM. n = 16-18 cells for each condition. An unpaired student’s *t*-test was performed (* p < 0.05).

The mobility of Munc18-1^WT^-mEos2 increased in stimulated DKO-PC12 cells compared to unstimulated cells (Fig. 5 B) as evidenced by the increased mobile population and concomitant reduction in the immobile fraction (Fig. 5 C-D). Although the distribution of the diffusion coefficients of Munc18-1^A297H^-mEos2 and Munc18-1^T304H^-mEos2 molecules in unstimulated DKO-PC12 cells also revealed two distinct populations (Fig. 5 F, I), there was no change in their mobility in response to stimulation (Fig. 5 E-J), suggesting that the transient binding of VAMP2 to Munc18-1 is essential for the release of Munc18-1 from its confined state.

The MSD and diffusion coefficient data was calculated from the average values of each analysed cell. These averaged values derived from >1000 molecules trajectories per cell are indicative of the overall changes in motion behavior. However, other analytical approaches are needed to infer whether Munc18-1 exhibit distinct motion states and whether these motion states and their transitions are affected by the mutations. We therefore applied Hidden Markov Modeling to our sptPALM-derived trajectories to determine the different mobility states ^46^ of the Munc18-1-mEos2 molecules. We found that Munc18-1 mobilities were grouped into four different diffusional states: S_1_: immobile, S_2_: slow diffusive, S_3_ and S_4_: fast diffusive (Fig. 6). When cells were stimulated to induce SV exocytosis, the apparent diffusion coefficients D_1_-D_4_ of the respective states did not change significantly (Fig. 6 A1, B1, C1), there was a significant decrease in the immobile occupancy state (S_1_) and an increase in the fast diffusive state although not significant (S_3_ and S_4_) (Fig. 6 A1-3). This switch in mobility state fits with the overall activity-dependent mobility increase observed in our sptPALM data (Fig. 5 B-D) and our cluster analysis showing smaller size clusters in response to stimulation (Fig. 4C). No significant changes in the state occupancies were observed for the VAMP2 binding-deficient Munc18-1^A297H^-mEos2 and Munc18-1^T304H^-mEos2 mutants (Fig 6 B, C). These results suggest that VAMP2-Munc18-1 interaction increases the probability of fast diffusive behavior, resulting in the release of a fraction of Munc18-1 molecules out of confinement.

**Figure 6:**
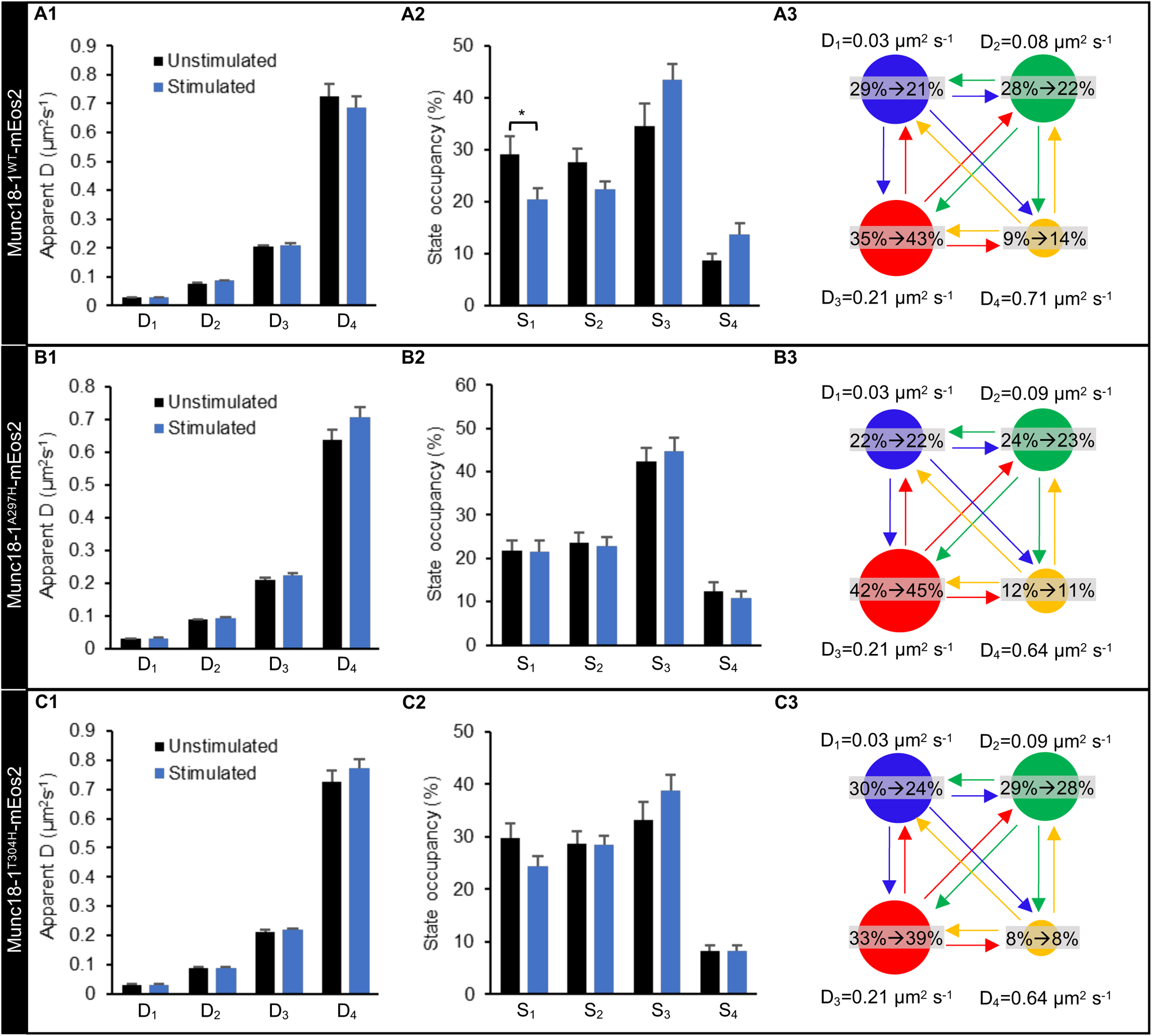
Binding to VAMP2 controls the activity-dependent increase in Munc18-1 diffusivity. Hidden Markov modeling with a four state model was used to determine the mobility states of **(A1-3)** Munc18-1^WT^-mEos2, **(B1-3)** Munc18-1^A297H^-mEos2 and **(C1-3)** Munc18-1^T304H^-mEos2 trajectories. **(A1, B1, C1)** Apparent diffusion coefficients of the four states (D_1_-D_4_), and **(A2, B2, C2)** respective state occupancy (S_1_-S_4_). Results are shown as mean ± SEM. An unpaired student’s *t*-test was performed (* p < 0.05). **(A3, B3, C3)** The four states (S_1_ immobile: blue; S_2_ slow diffusive: green; S_3_ and S_4_ fast diffusive: red and yellow respectively) are presented as colored circles, the areas of which are proportional to the respective state occupancy.

### The activity-dependent mobility decrease of syntaxin-1A is dependent upon the interaction of Munc18-1 with VAMP2

We have previously shown that the Munc18-1 domain 3a hinge-loop controls the activity-dependent confinement of syntaxin-1A in nanodomains during SNARE complex assembly ^25^. To assess the effect of VAMP2 binding to Munc18-1 on the mobility of syntaxin-1A, we performed sptPALM of syntaxin-1A-mEos2 expressed in DKO-PC12 cells co-transfected with either Munc18-1^WT^-emGFP or with the VAMP2 binding-deficient mutants Munc18-1^A297H^-emGFP or Munc18-1^T304H^-emGFP. A low resolution TIRF image shows a DKO-PC12 cell expressing syntaxin-1A-mEos2 and Munc18-1^WT^-emGFP with a non-uniform distribution across the plasma membrane (Fig. 7 A1). The super-resolved sptPALM-intensity, trajectory and diffusion coefficient maps showed that syntaxin-1A-mEos2 is organized in nanoclusters across the whole plasma membrane as previously reported (Fig. 7 A2-A4) ^30,47^. The quantitative analysis of syntaxin-1A-mEos2 trajectories revealed a significant decrease in syntaxin-1A-mEos2 mobility after stimulation in the DKO-PC12 cells rescued with Munc18-1^WT^-emGFP (Fig. 7 B). Interestingly, this activity-dependent decrease in syntaxin-1A-mEos2 mobility was lost in DKO-PC12 cells expressing the VAMP2 binding-deficient mutant Munc18-1^A297H^-emGFP (Fig. 7 C, E). In cells expressing Munc18-1^T304H^-emGFP, the mobility of syntaxin-1A-mEos2 decreased after stimulation (Fig. 7 D), however this effect was not as pronounced as in cells expressing Munc18-1^WT^-emGFP (Fig. 7 B, E). This indicates that the interaction of VAMP2 with the Munc18-1 domain 3a A297 residue and to a lesser extent T304, is essential for the activity-dependent confinement of syntaxin-1A.

**Figure 7:**
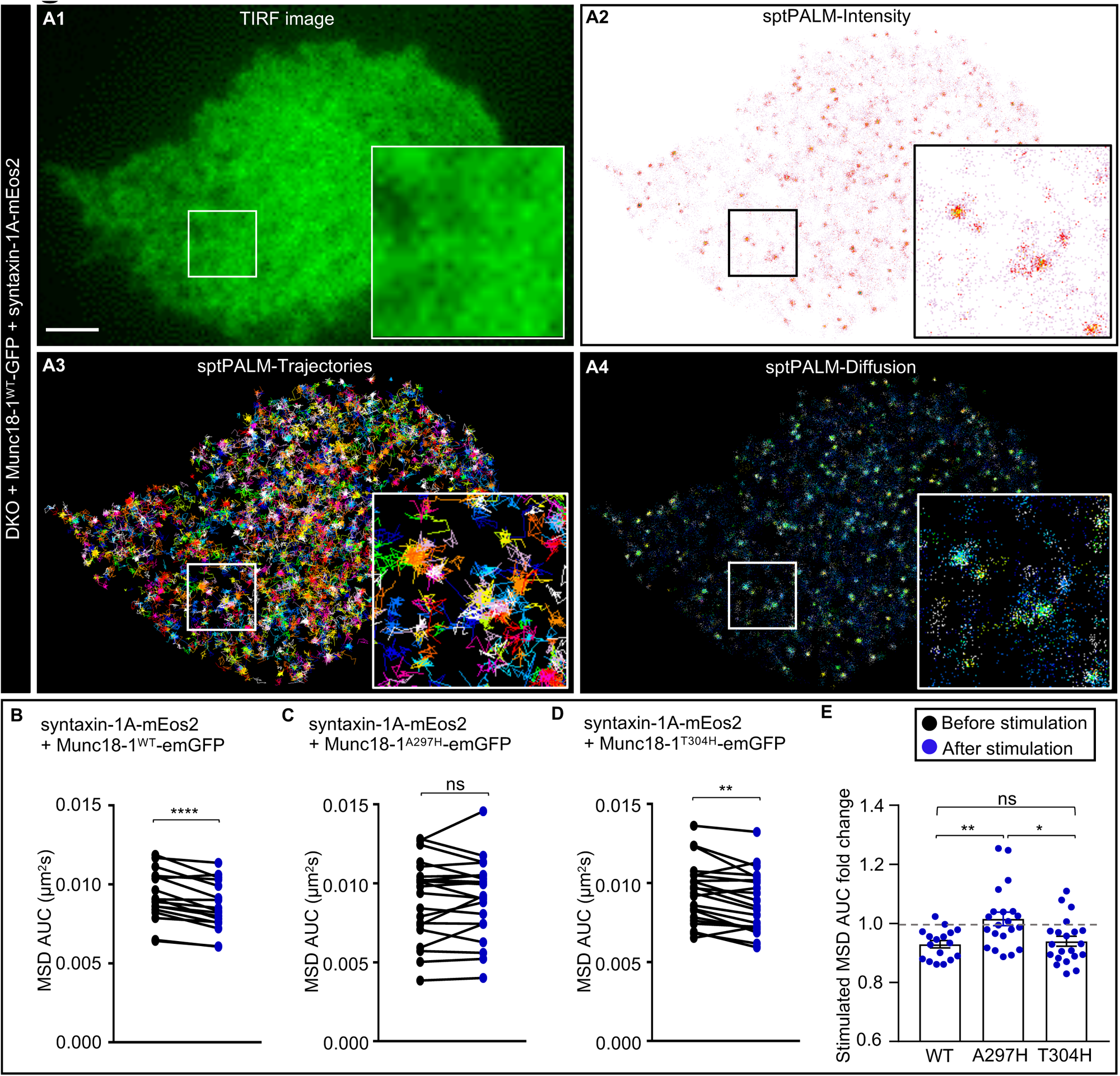
Munc18-1 binding to VAMP2 controls the activity-dependent mobility decrease of syntaxin-1A following SNARE complex formation. DKO-PC12 cells co-transfected with syntaxin-1A-mEos2 and either **(A-B)** Munc18-1^WT^-emGFP, (**C)** Munc18-1^A297H^-emGFP or **(D)** Munc18-1^T304H^-emGFP were imaged at 50 Hz before and after stimulation with 2 mM BaCl_2_. Representative **(A1)** TIRF image, and corresponding **(A2)** sptPALM average intensity map, **(A3)** a trajectory map (trajectory color-coding in arbitrary units), **(A4)** diffusion coefficient map, of a DKO-PC12 cell co-expressing syntaxin-1A-mEos2 and Munc18-1^WT^-emGFP (scale bar: 2 µm). Enlarged inserts from the region of interest are shown in the respective panels. The average area under the curve (AUC) of the mean square displacement (MSD) is shown for syntaxin-1A-mEos2 in DKO-PC12 cells expressing either **(B)** Munc18-1^WT^-emGFP, **(C)** Munc18-1^A297H^-emGFP or **(D)** Munc18-1^T304H^-emGFP, before (black) and after (blue) stimulation (n = 17-21 cells for each condition). **(E)** MSD AUC fold change after stimulation. Results in **(E)** are expressed as mean ± SEM. (B-D) A paired student’s independent *t*-test, and **(E)** one-way ANOVA with multiple comparisons was performed (ns, not significant; * p < 0.05; ** p < 0.01; **** p < 0.0001).

## Discussion

Our study endeavored to characterize the nanoscale molecular steps enabling the formation of a fusion-competent interface between the plasma membrane and SVs. We had previously shown that the release of Munc18-1 from nanoclusters is an essential step that allows syntaxin-1A to open and engage in SNARE complex assembly. However, it is not clear whether the opening of syntaxin-1A and release of Munc18-1 occurs before or as a result of vesicular docking. The results presented here suggest that Munc18-1 could catalyze SNARE complex assembly upon docking of the vesicle via opening of syntaxin-1A within the context of SNAP25 and VAMP2. As SNAP25 has been shown to be preassembled with syntaxin-1A/Munc18-1 on the plasma membrane ^43,48,49^, we propose that the transient binding of VAMP2 to Munc18-1 domain 3a triggers the opening of syntaxin-1A, initiating its engagement into the SNARE complex which triggers the subsequent release of Munc18-1.

Our data demonstrates that the A297H (and to a lesser extent T304H) mutation impairs the fusion of SVs resulting in prolonged docking of SVs on the plasma membrane of DKO-PC12 cells creating a ‘super-docking’ phenotype. A similar super-docking phenotype has been previously reported for a Munc18-1 domain 3a hinge-loop (317-333) deletion mutant ^25^. This could either suggest that the VAMP2 binding deficient mutants affect the extension of domain 3a, thereby phenocopying the effect of the hinge-loop deletion previously reported, or that the binding of VAMP2 to Munc18-1 might be required for the full zippering of the SNARE complex and subsequent release of Munc18-1. A significant reduction in SV fusion was observed for the Munc18-1^A297H^ mutant, suggesting that vesicles could be locked in a docked state as a result of an arrested priming step, rendering them unable to undergo complete SNARE zippering. Although the structure does not indicate precisely why there is a difference in neuroexocytosis impairment caused by the Munc18-1^A297H^ and Munc18-1^T304H^ mutants, the stronger effect indicates that the A297H mutation poses a stronger steric hindrance than the T304H mutation ^33^. Other *in vitro* studies have provided evidence that Munc18-1 can form a template for SNARE complex assembly ^28,37,50^, and that it binds to synaptobrevin (VAMP2) using biophysical approaches ^51^. This is in agreement with our results presented here, which confirm that Munc18-1 must form a template for SNARE complex assembly *in vivo* in neurosecretory cells. It has also been suggested that SNARE complexes are in equilibrium between half- and full-zippered states^52^. The binding of VAMP2 at the A297 residue of Munc18-1 might be sufficient to shift the equilibrium towards fully-zippered states which could be the reason why the T304H mutation has a less pronounced effect than A297H on exocytosis.

Our results also demonstrate that the binding of VAMP2 to Munc18-1 is required for the activity-dependent release of Munc18-1^WT^-mEos2 from nanoclusters on the plasma membrane. When the binding to VAMP2 is impaired, Munc18-1 seems to be stuck in nonfunctional nanoclusters, likely in aborted complexes with syntaxin-1A. In addition, the activity-induced mobility increase of Munc18-1^WT^-mEos2 is lost in Munc18-1^A297H^-mEos2- and Munc18-1^T304H^-mEos2-expressing cells. This indicates that the binding of VAMP2 to Munc18-1 controls the dynamic reorganization of Munc18-1 nanoclusters, which provides a signature for successful vesicular priming and is critical for SV fusion. Moreover, the activity-dependent trapping of syntaxin-1A was lost in the presence of the VAMP2 binding-deficient Munc18-1^A297H^-mEos2 mutant. Thus, the interaction of VAMP2 with Munc18-1 also controls the dynamic nanoscale organization of syntaxin-1A which is required for the engagement of syntaxin-1A into SNARE complexes ^25^. This supports the hypothesis that VAMP2 binding to Munc18-1 domain 3a initiates the opening of syntaxin-1A by promoting the extension of the domain 3A hinge-loop, which in turn allows the templating and engagement of VAMP2 and syntaxin-1A into the SNARE complex assembly. In addition, these results show that the interaction of Munc18-1 with VAMP2 not only controls SNARE assembly, but also the dynamic nanoscale organization of Munc18-1 and syntaxin-1A and possibly of other proteins involved in SV priming and fusion.

Munc18-1, together with Munc13-1, promotes the formation of the SNARE complex, competing against NSF/αSNAP-dependent SNARE unzippering ^14,15^. It is still not clear if the release of Munc18-1 has a role in the events following SV fusion. Munc18-1 could also contribute to other priming interactions involving Munc13-1 and αSNAP ^15,53^ most likely while Munc18-1 is still in complex with syntaxin-1A and VAMP2. It has been shown that Sec/Munc18 proteins regulate SNARE complex formation by selectively binding to SNARE molecules and protecting them from disassembly by NSF and αSNAP ^15,54,55^. Thus in addition to being a master regulator of priming, Munc18-1 also plays a key role in preventing de-priming by NSF and αSNAP ^54,55^.

We propose a model for the role for the interaction of VAMP2 with the Munc18-1 domain 3a during SNARE complex formation (Fig. 8). Munc18-1 is in an inhibitory state while in complex with closed syntaxin-1A. Upon stimulation the Munc18-1 domain 3a loop changes to an extended conformation which mediates the opening of syntaxin-1A. This conformational change of the Munc18-1 domain 3a loop is likely initiated through an interaction with VAMP2 on docked vesicles. This step may stabilize Munc18-1 domain 3a in the extended hinge-loop conformation, facilitating the opening and engagement of syntaxin-1A with the cognate SNARE proteins. This could allow the SNAREs to form functional partial zippers orienting from the N-terminal to the C-terminal region ^1,56,57^. Upon successful zippering of the SNAREs, Munc18-1 is released from the SV fusion site. Other exocytic molecules are also involved in priming the SV-plasma membrane interface.

**Figure 8:**
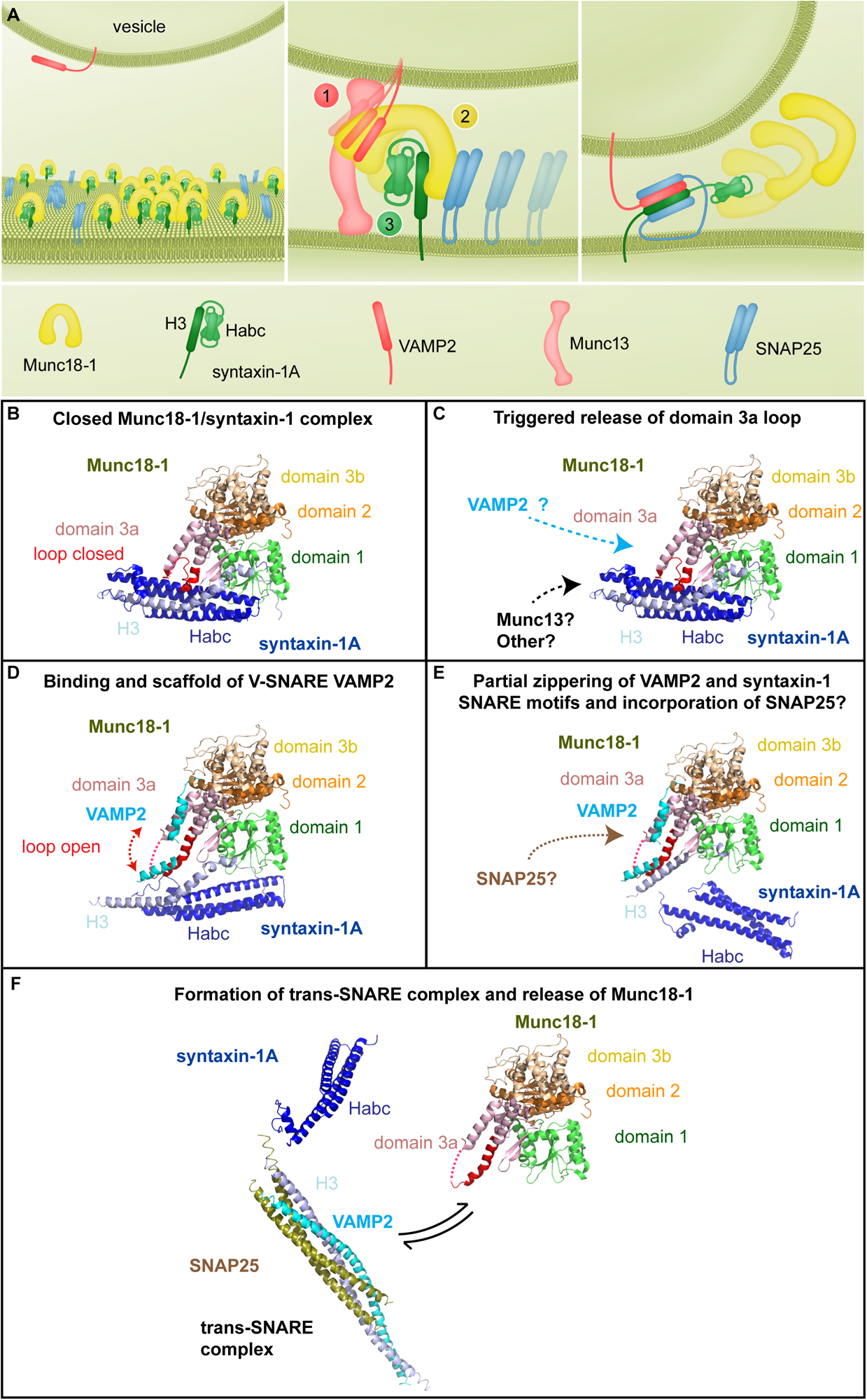
Schematic of the SNARE proteins during Munc18-1 domain 3a conformational switch. **(A)** Cartoons showing the dynamic molecular interactions preceding SNARE complex formation. The first panel shows the nanodomain organization of syntaxin-1A (green) in complex with Munc18-1 (yellow) and SNAP25 (blue) on the plasma membrane, and VAMP2 (red) on the vesicle. The second panel illustrates transient VAMP2 binding to the (1) extending Munc18-1 domain 3a (2), favoring its extension which in turn relaxes syntaxin-1A allowing it to open (3). The last panel shows the exit of Munc18-1 from the SNARE complex. **(B)** Syntaxin-1A interacts with Munc18-1 in a closed conformation during trafficking, and prior to the stimulation of exocytosis. **(C)** Possible binding of VAMP2 and other proteins with Munc18-1. **(D)** A conformational change in the Munc18-1 domain 3a loop (triggered by VAMP2 binding) allows syntaxin-1A to open. **(E-F)** The domain 3a loop provides a template for interaction with the v-SNARE VAMP2, allowing assembly of syntaxin-1A with VAMP2 and possibly SNAP25 to integrate into the intermediary SNARE complex. **(F)** Release of Munc18-1 from the SNARE complex assembly.

The weak interaction of Munc13-1 with syntaxin-1A orchestrates the opening and SNARE complex engagement of syntaxin-1A together with Munc18-1 ^14–16^. Munc18-1 domain 3a conformational change and Munc13-1 interaction may occur simultaneously, leading to an efficient priming. Our binding experiments in cells, suggest that other priming proteins could also regulate the binding of Munc18-1 to VAMP2. Indeed, Munc13-1 was recently shown to trigger a conformational change in the Munc18-1 domain 3a from an autoinhibitory state to an extension state which allows VAMP2 binding ^37^, and the Munc18-1^P335A^ mutant whose binding to VAMP2 is increased, bypasses the requirement for Munc13-1 ^28,58^. In this view, a recent study showed that mutations T323A/M324A/R325A within the Munc18-1 domain 3a inhibit SNARE complex assembly in a Munc13-1-dependent manner, when Munc18-1 was precomplexed with syntaxin-1A. These mutations did not hinder Munc18-1’s ability to either bind to syntaxin-1A or VAMP2^59^. Pertsinidis et al. ^60^ showed that Munc18-1 is associated with the SNARE protein SNAP25 in complex and that Munc18-1 nanoclusters on the plasma membrane colocalize with syntaxin-1A and SNAP25 nanoclusters, indicating that Munc18-1 is not released when SNAP25 engages into the SNARE complex, but potentially before the full zippering of the complex. This is consistent with recent results showing that Munc13-1 and Munc18-1 serve as a template for a half zippered SNARE assembly, from which Munc18-1 dissociates to allow full zippering ^61^. Our study further showed that alterations in the ability of Munc18-1 to bind to VAMP2 impacts the nanoscale mobility of Munc18-1, its clustering behavior on the membrane, and its ability to reorganize upon stimulation. The precise control of Munc18-1 domain 3a engagement into SNARE complex at the SV-plasma membrane interface during fusion is therefore dependent upon its ability to bind and facilitate effectively zippering of VAMP2 with other v-SNAREs.

## Acknowledgements

We thank Jean-Baptiste Sibarita (University of Bordeaux) for providing the single-molecule analysis software PalmTracer, Rumelo Amor and all past and present members of the Microscopy and Microanalysis facility at the Queensland Brain Institute (QBI) for their outstanding microscopy support and assistance in super-resolution analysis, as well as Jake Carrol and the QBI IT department. We also thank Nick Valmas for drawing the schematic model, Alex McCann and Rowan Tweedale for critical reading of this article. In addition, we thank Prof Jennifer Martin for co-supervision of E.K.L. and guidance in this project. This work was supported by a Discovery Early Career Research Award (DE190100565 to M.J.), an Australian Research Council (ARC) Discovery Project (DP170100125 to F.A.M.), a National Health and Medical Research Council (NHMRC) Project Grant (APP1138083 to B.M.C. and F.A.M.), and an ARC Linkage Infrastructure, Equipment and Facilities LIEF Grant (LE130100078 to F.A.M.). F.A.M. is an NHMRC Senior Research Fellow (GNT1060075) and B.M.C. is holds an NHMRC Investigator Grant (APP2016410). E.K.L. is supported by a UQ Research Scholarship, an IMB Research Advancement Award, and an AINSE Postgraduate Research Award. We also acknowledge the use of the Australian Microscopy & Microanalysis Research Facility at the Centre for Microscopy and Microanalysis at The University of Queensland.

## Author Contributions

B.M.C. performed the structural modelling and identified the Munc18-1 residues, which were mutated for this project. R.K. performed and analysed the SV tracking and Munc18-1 sptPALM experiments. R.K and M.M. performed the Munc18-1 fixed PALM experiments and T.P.W. analysed the data by DBSCAN. T.P.W also developed the BOOSH analysis pipeline. A.B. and J.Z.B. performed the syntaxin-1A sptPALM experiment and A.B. analysed the data. M.S. performed and analysed the co-immunoprecipitation experiments in DKO-PC12 cells. Y.J.C. generated the DKO-PC12 cells. A.J. performed and analysed the VAMP2-pHluorin exocytosis assay, which was supervised by M.J. and R.M.. S.K.W. and E.K.L. performed and analysed the Munc18-1 and VAMP2-GST pulldowns, which were designed and supervised by B.M.C.. R.S.G. performed and analysed the NPY-hPLAP release assays for the DKO-PC12 cells, the syntaxin-1A immunofluorescence staining, mRNA sequencing, and BOOSH cluster analysis. P.P. performed the Hidden Markov modelling. A.B., R.K., R.S.G., B.M.C. and F.A.M. wrote the manuscript. All co-authors contributed to the critical appraisal of the manuscript. F.A.M. designed and supervised the study.

## Supplementary Figure Legends

**Figure S1:**
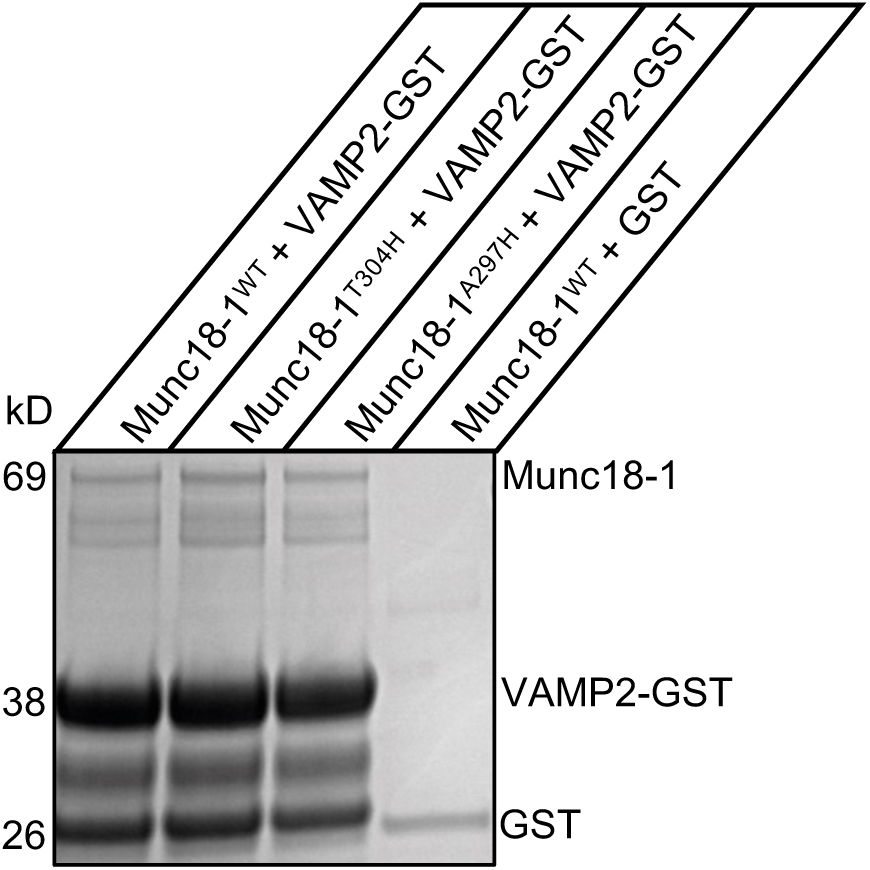
The VAMP2/Munc18-1 interaction is transient and relies on the Munc18-1 A297 residue. Full version of the Coomassie blot shown in Figure 1B: Recombinantly expressed Munc18-1^WT^-His, Munc18-1^T304H^-His or Munc18-1^A297H^-His was incubated with VAMP2-GST and glutathione agarose beads overnight at pH 8.0, immobilized on glutathione agarose resin, analyzed by SDS-PAGE and the gels stained with Coomassie blue. As a control, Munc18-1^WT^-His was also incubated with GST alone. The results were reproduced in three independent experiments.

**Figure S2:**
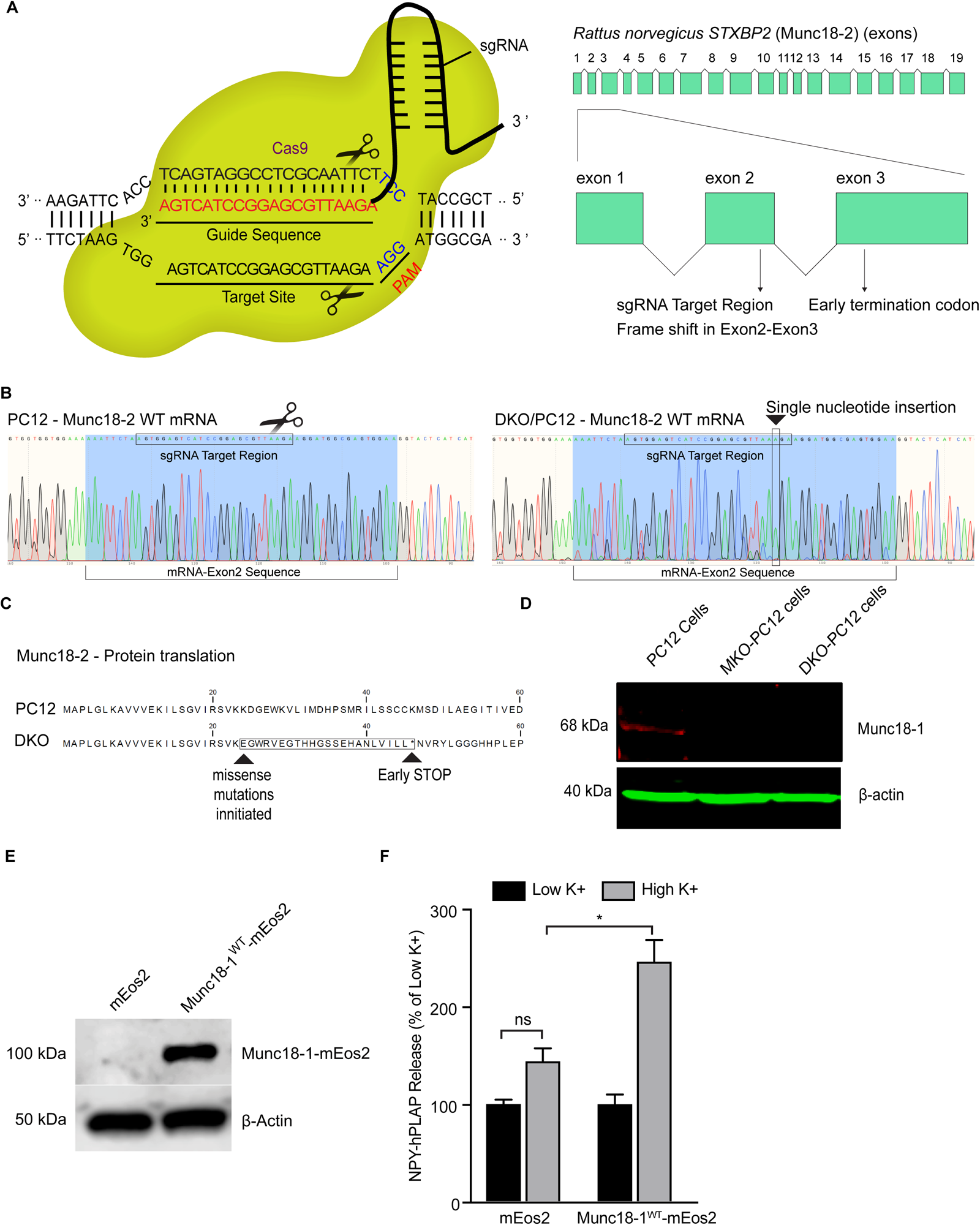
Generation and validation of Munc18-1/2 double knockout PC12 cells (DKO-PC12). **(A)** Schematic diagram of the sgRNA target site (Munc18-2 exon2). **(B)** Sequence chromatogram of mRNA from PC12 cells (left) or double knock-out Munc18-1/2 (DKO)-PC12 cells (right). Note the single-nucleotide insertion into exon 2 of Munc18-2 in the Munc18-1/2 DKO-PC12 cells (indicated with the black arrow head). (**C**) Translated Munc18-2 protein sequence in DKO-PC12 cells as a result of the frame shift in exon2, results in an early stop codon. **(D)** Cell lysates of wild-type PC12 cells, MKO-PC12 cells and clonal DKO-PC12 cells were subjected to western blotting analysis and probed with anti-Munc18-1 antibody. A parallel set of lysates were probed with anti-actin antibody as a loading control. **(E)** DKO-PC12 cells were co-transfected with NPY-hPLAP and either empty pmEos2-N1 vector or Munc18-1^WT^-mEos2. Transfected cells were analysed by western blotting. Note that DKO-PC12 cells expressing pmEos2-N1 alone do not contain any endogenous Munc18-1 (∼70 kDa). β-Actin was used as a loading control. **(F)** Cells were stimulated with 2 mM BaCl_2_ to elicit secretion for 5 min. Released NPY-hPLAP was expressed as a percentage of release in unstimulated cells. Results are expressed as mean ± SEM. One-way ANOVA with multiple comparisons. * = p < 0.05.

**Figure S3:**
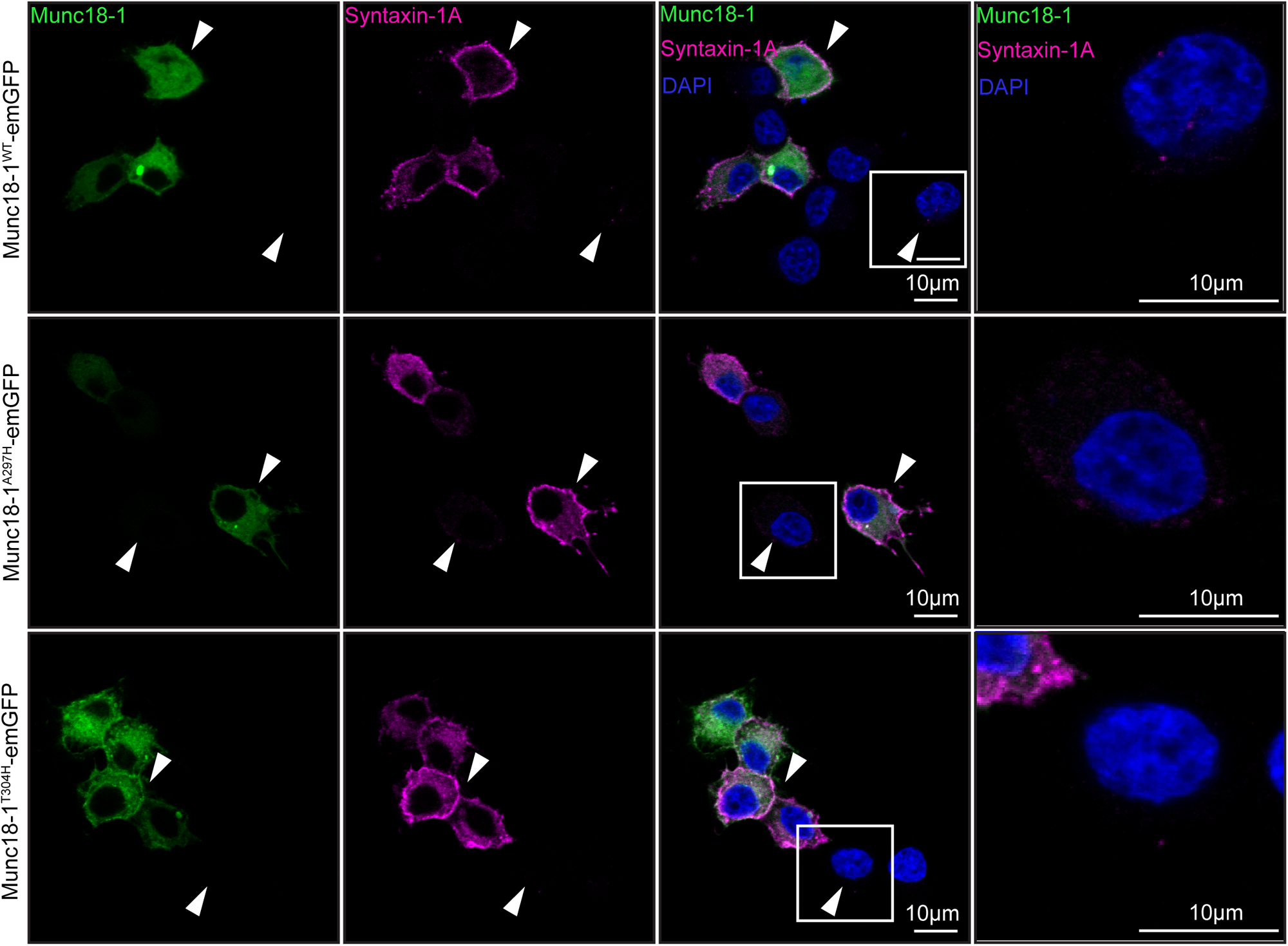
VAMP2 binding-deficient Munc18-1 mutants rescue syntaxin-1A localization on the plasma membrane. DKO-PC12 cells were co-transfected with syntaxin-1A-mEos2 and either Munc18-1^WT^-emGFP, Munc18-1^A297H^-emGFP or Munc18-1^T304H^-emGFP. Cells were then fixed and labelled for syntaxin-1A (anti-syntaxin/Alexa647) and DAPI, then mounted and imaged using confocal microscopy. Representative images show (L-R) Munc18-1 (green), syntaxin-1A (magenta) and DAPI (blue in merge). Arrowheads pointing down indicate cells with syntaxin-1A plasma membrane labelling, arrowheads pointing up indicate non-transfected cells which do not express Munc18-1 and show low endogenous syntaxin-1A expression and plasma membrane localization (scale bars: 10 μm). Far right panels – zoom of untransfected cells highlighting transport deficit of syntaxin-1A in the absence of Munc18-1/2.

**Figure S4:**
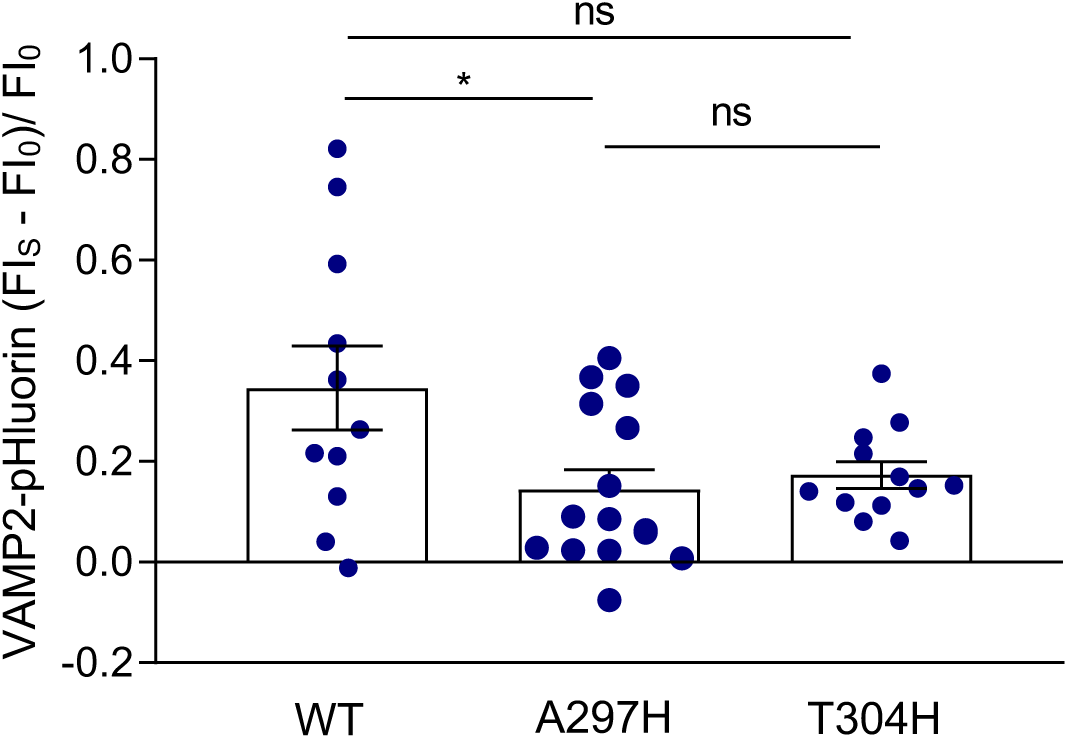
Secretory vesicle exocytosis is decreased in VAMP2 binding-deficient Munc18-1 mutants. DKO-PC12 cells were co-transfected with VAMP2-pHluorin and either Munc18-1^WT^-mCherry, Munc18-1^A297H^-mCherry, or Munc18-1^T304H^-mCherry. SV release was determined by the fluorescence intensity (FI) fold change of VAMP2-pHluorin at 200 s after high K^+^ stimulation. Results are expressed as mean ± SEM. A Kruskal-Wallis test with multiple comparisons was performed (ns, not significant; * p < 0.01). n=11 for Munc18-1^WT^-mCherry, n=15 for Munc18-1^A297H^-mCherry, n=12 for Munc18-1^T304H^-mCherry.

**Figure S5:**
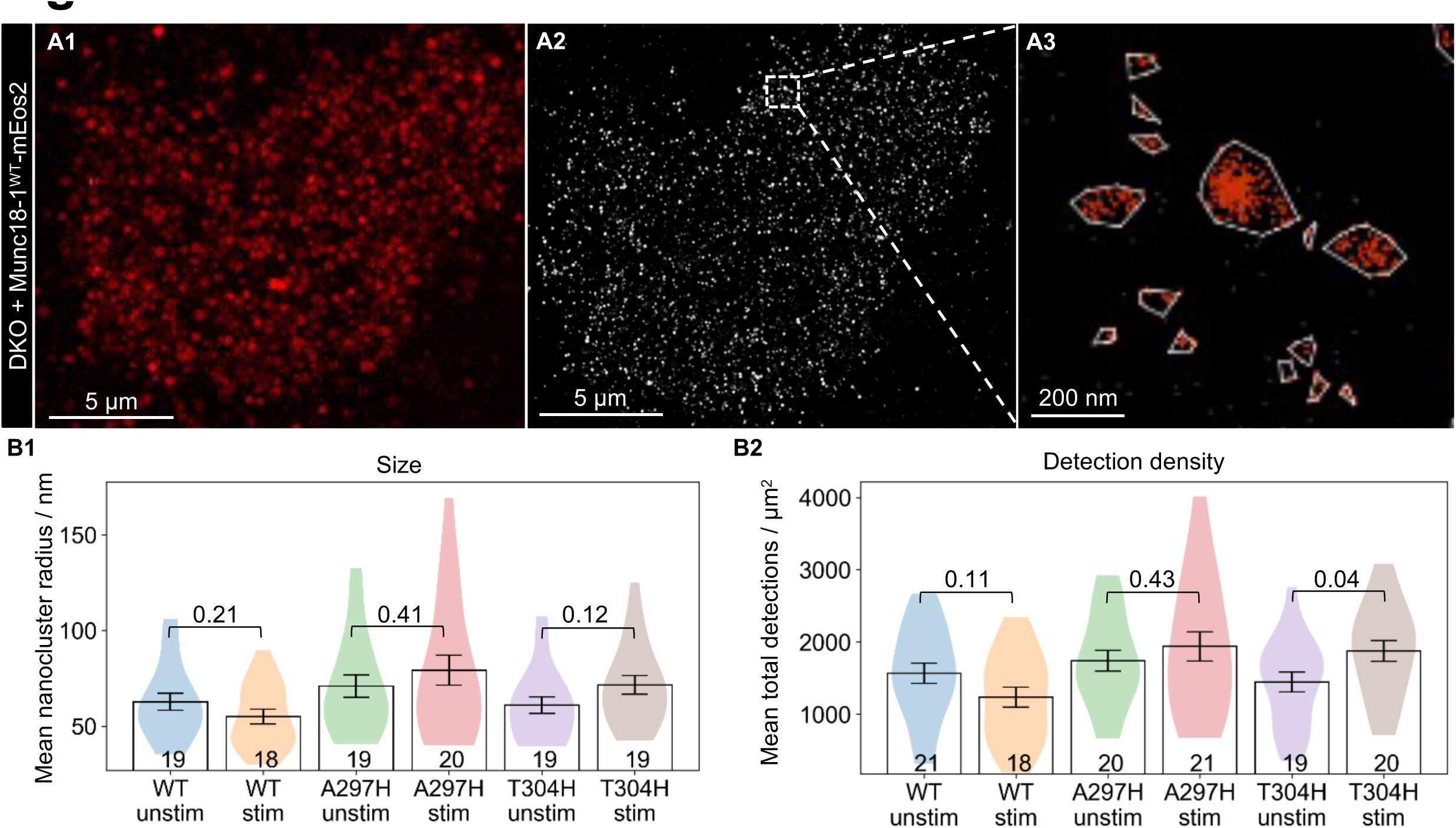
Munc18-1 binding to VAMP2 controls the activity-dependent release of Munc18-1 from nanoclusters on the plasma membrane. DKO-PC12 cells expressing either Munc18-1^WT^-mEos2, Munc18-1^A297H^-mEos2 or Munc18-1^T304H^-mEos2 were stimulated for 2 min with 2 mM BaCl_2_ before fixation and were subjected to PALM in TIRF. **(A1)** Maximum intensity projection of a 20000 frame movie of a DKO-PC12 cell expressing Munc18-1^WT^-mEos2. **(A2)** The super resolved PALM image and **(A3)** magnified view of (A2). **(B)** DBSCAN and custom Python scripts were used to quantify the **(B1)** size (mean nanocluster radius/nm), **(B2)** detection density (mean total detections/µm^s^). An unpaired student’s *t*-test was performed and p values are shown (p <0.05 was considered significant).

## STAR Methods

### CONTACT FOR REAGENT AND RESOURCE SHARING

Further information and requests for resources and reagents should be directed to and will be fulfilled by the Lead Contact, Frédéric A. Meunier. All reagents and resources used in this paper are subject to MTAs.

### EXPERIMENTAL MODEL AND SUBJECT DETAILS

#### Cell line

DKO-PC12 cells were maintained at 37 °C, 5 % CO_2_ in Dulbecco’s modified Eagle’s medium (DMEM) (Thermo Fisher Scientific) supplemented with 5 % fetal calf serum (Bovogen), 5 % heat-inactivated horse serum (Gibco, Invitrogen) and 0.5 % GlutaMAX™ (Gibco, Thermo Fisher Scientific). Cells were transfected using the Lipofectamine™ LTX (Thermo Fisher Scientific) reagent according to the manufacturer’s instructions. For imaging, the cells were plated on poly-D-lysine-coated glass-bottom culture dishes (Cellvis) after 48 h.

#### *E. coli* strains used for protein expression

*E. coli* Rosetta cells were cultured at 37 °C in lysogeny broth (LB) (tryptone 10 g/L; yeast extract 5 g/L; NaCl 5 g/L) shaking at 180 rpm. When the cells reached an optical density (OD_600_) of 0.8, protein overexpression was induced with 0.5 mM isopropyl β-D-1-thiogalactopyranoside (IPTG) and the cells were grown at 20 °C shaking at 180 rpm overnight.

*E. coli* BL21(DE3) cells (New England Biolabs) were cultured at 37 °C in lysogeny broth (LB) shaking at 180 rpm. When the cells reached an optical density (OD_600_) of 0.5-0.7, protein expression was induced with 0.5 mM isopropyl β-D-1-thiogalactopyranoside (IPTG) and the cells were grown at 23 °C shaking at 180 rpm overnight.

### METHOD DETAILS

#### Plasmids

Munc18-1^A297H^ and Munc18-1^T304H^ constructs were generated using the QuickChange Lightning Site-Directed Mutagenesis Kit (Agilent Technologies) to generate point mutations. Fluorescently labelled Munc18-1^A297H^ constructs were amplified by PCR using the following primers: forward, 5’-GCGACACAAGCACATCCACGAGGTGTCCCAGGAAG-3’; and reverse, 5’-CTTCCTGGGACACCTCGTGGATGTGCTTGTGTCGC-3’, and fluorescently labelled Munc18-1^T304^ constructs were amplified by PCR using the following primers: forward; 5’-CAGAGGTGTCCCAGGAAGTGCACCGGTCTCTGAAGGACTTTTC-3’; and reverse, 5’-GAAAAGTCCTTCAGAGACCGGTGCACTTCCTGGGACACCTCTG-3’. Munc18-1^WT^-mEos2 was used as a PCR template for Munc18-1^A297H^-mEos2 and Munc18-1^T304H^-mEos2; Munc18-1^WT^-mCherry for Munc18-1^A297H^-mCherry and Munc18-1^T304H^-mCherry, and Munc18-1^WT^-emGFP for Munc18-1^A297H^-emGFP and Munc18-1^T304H^-emGFP. GST-tagged and His-tagged Munc18-1^A297H^ were amplified by PCR using the following primers: forward, 5’ CTGCGACACAAGCACATCCACGAGGTGTCCCAGGAAGTG 3’; and reverse: 5’ CACTTCCTGGGACACCTCGTGGATGTGCTTGTGTCGCAG 3’, and GST-tagged and His-tagged Munc18-1^T304H^ were amplified by PCR using the following primers: forward, 5’ GCAGAGGTGTCCCAGGAAGTGCACCGGTCTCTGAAGGACTTTTC 3’; reverse, 5’ GAAAAGTCCTTCAGAGACCGGTGCACTTCCTGGGACACCTCTGC 3’.

Munc18-1-His in a pEt28a vector and GST-Munc18-1 in a pGEX4T-2 vector (see below) were used as PCR templates respectively.

A VAMP2_1-94_ with a non-cleavable C-terminal GST tag was cloned by amplifying two PCR products using the following primers: forward, 5’ AAGGAGATATACATATGTCGGCTACCGCTGCCACCGTC 3’; reverse, 5’ CTAGTATAGGGGAGCCCTTGAGGTTTTTCCACCAGTATTTGCGCTTGAGCTTGG 3’ and forward, 5’ GTGGAAAAACCTCAAGGGCTCCCCTATACTAGGTAAATGGAAAATTAAGGGCCTT 3’; reverse, 5’ GACGGAGCTCGAATTCTTATTAGGTACCGCCGCCACCACTAGTATCCGATTTTGGAG 3’ using GST-VAMP2 in a pGEX4T-2 vector as a template. The products from the previous two PCRs were then used as template with the following primers: forward, 5’ AAGGAGATATACATATGTCGGCTACCGCTGCCACCGTC 3’; reverse, 5’ GACGGAGCTCGAATTCTTATTAGGTACCGCCGCCACCACTAGTATCCGATTTTGGAG 3’. This PCR product was cloned into pET24a linearised with Nde1 and EcoR1 using the standard procedure for the InFusion HD Cloning Kit (Clontech) to generate VAMP2-GST. All constructs were sequenced in the Australian Genome Research Facility at the University of Queensland.

pCMV-neuropeptide Y-emGFP (NPY-emGFP) was provided by S. Sugita (University of Toronto and University Health Network, Toronto, Canada).

Munc18-1-mEos2 and syntaxin-1A-mEos2 were generated as described previously ^25^. VAMP2-pHluorin was provided by J. Rothman (Yale University, New Haven, CT)^41^.

Munc18-1-His and syntaxin-1A-His in a pEt28a vector and GST-Munc18-1, GST-SNAP25 and GST-VAMP2 in a pGEX4T-2 vector were provided by Prof J. Martin (Griffith University, Brisbane, Australia).

#### GST-VAMP2 pulldown assays

Munc18-1^WT^-His, Munc18-1^A297H^-His, Munc18-1^T304H^-His and GST-VAMP2 were recombinantly expressed and purified from *E. coli* BL21(DE3) cells. Glutathione agarose beads (50 µl) were washed with 3 x 1 ml pulldown buffer (25 mM HEPES pH 7.5, 300 mM NaCl, 10 % (v/v) glycerol, 1 % (v/v) Triton X-100, 2 mM β-mercaptoethanol). Munc18-1^WT^ and mutant proteins (1 nmol) were combined with either GST-VAMP2 (0.5 nmol) or GST (0.5 nmol) in pulldown buffer (total volume 250 µl) and centrifuged at 17,000 rpm for 10 min before the addition of the beads. The samples were incubated at 4 °C on a rocker overnight. The beads were then washed with 3 x 1 ml of pulldown buffer. Samples were run on a 10 % Bis-Tris SDS PAGE gel in MES buffer at 165 V for 35 min, then stained with Coomassie blue. The experiment was repeated three times.

#### GST-Munc18-1 pulldown assays

Pulldown assays were carried out using GST-tagged Munc18-1^WT^, as well as GST-tagged Munc18-1^A297H^ and Munc18-1^T304H^ mutants. GST-Munc18-1^WT^, GST-Munc18-1^A297H^, GST-Munc18-1^T304H^, syntaxin-1A-His, GST-SNAP25 and GST-VAMP2 were recombinantly expressed and purified from *E. coli* Rosetta cells. The GST tags of GST-SNAP25 and GST-VAMP2 were cleaved off using the enzyme Thrombin with an overnight incubation at 4 °C. The SNARE complex was formed by incubating syntaxin-1A-His with an excess of SNAP25 and VAMP2 overnight at 4 °C, after which 1 nmol of the pre-formed ternary complex was incubated with 0.5 nmol GST-tagged Munc18-1^WT^ or mutants at 4 °C in 500 µl of pulldown buffer (50 mM Tris-HCl pH 8, 200 mM NaCl, 0.1 % Triton X-100 and 2 mM β-mercaptoethanol). 50 µl of glutathione Sepharose beads were then added to the protein mix and incubated for 30 min at 4 °C, after which the beads were washed thoroughly using the above buffer and bound proteins were analyzed by reducing SDS-PAGE, with Coomassie blue staining. This experiment was performed twice.

#### Generation of the DKO-PC12 cell line

Munc18-1 knockout PC12 (MKO-PC12) cells ^62^ were cultured in 6-well dishes to 70-80 % confluence. Cells were transfected with 1 μg sequence verified pSpCas9-Munc18-2 (sgRNA)-2A-GFP using Lipofectamine 2000 (Invitrogen). For double knockout Munc18-1/2 PC12 cells (DKO-PC12 cells), a 20 bp guide sequence (5’-AGTCATCCGGAGCGTTAAGA-3’) targeting DNA within the second exon of Munc18-2 was selected from an online CRISPR design tool. The sgRNA expression construction method has been described previously ^63^. 48 h post transfection, cells were pelleted in Dulbecco’s modified eagle medium (DMEM) and sorted in 96 well plates using a FACSAria II cell sorter (BD Biosciences). Single cells from GFP-expressing cells (high expression population) were expanded to obtain individual clones.

Individual clones were lyzed with ice-cold cell lysis buffer (1 % Triton X-100, 1 mM EDTA, 1 mM EGTA, and 5 mM Na-pyrophosphate in PBS) supplemented with protease inhibitor cocktails and centrifuged at 14,000 rpm for 20 min at 4 °C. The whole cell lysates were resuspended in 2X SDS sample buffer and analyzed by western blot with a mouse anti-Munc18-1 antibody (BD Biosciences, 610336). Genomic DNA was isolated from edited clones and non-edited PC12 cells (control). A region of exon2 of the Munc18-2 gene was amplified with genomic DNA specific primers (forward primer, 5’-CGGAGTCCGCGCGTCAGTCGGT-3’; reverse primer, 5’-ATAAAGGGGCGGATGGGGGAGGGA-3’). The PCR products were A-tailed and cloned into pGEM-T easy (Promega) and sequenced in the Australian Genome Research Facility at The University of Queensland. RNA from PC12 and DKO-PC12 cells was extracted using RNA isolation kit (Machrey-Nagel) and used to generate PolyA-tailed mRNA by RT-PCR using dT23VN primer. Forward Primers (binding to Exon1 of Munc18-2) F’ ‘GCGGTGGTGGTGGAAAAA’ and reverse primer (exon-exon spanning) ‘ACCACATCCTCCATCACGTC’ to produce a 1,440 bp amplicon (confirmed by agarose gel electrophoresis, data not shown). Amplicons were purified by PCR clean up and send for sanger sequencing (AGRF) with reverse primer binding to exon 4 of Munc18-2 M18-2 Ex4Seq Rev primer ‘AAGCTGGGAATGGGTTCTCT’.

#### NPY-hPLAP release assays

DKO-PC12 cells were co-transfected with Neuropeptide Y (NPY) fused to the catalytic domain of human placental alkaline phosphatase (NPY-hPLAP) and either pmEos2-N1 control plasmid or Munc18-1^WT^-mEos2 for 48 h. Cells were washed and incubated with buffer A as a control (145 mM NaCl, 5 mM KCl, 1.2 mM Na_2_HPO_4_, 10 mM D-glucose, 20 mM Hepes, pH 7.4) or stimulated with 2 mM BaCl_2_ for 5 min at 37 °C. NPY-hPLAP released from cells was measured using the chemiluminescent reporter gene assay system (Phospha-Light; Applied Biosystems) according to the manufacturer’s instructions, and the results were expressed as a percentage of control.

#### Immunoprecipitation

For immunoprecipitation, DKO-PC12 cells were transfected with Munc18-1^WT^-emGFP, Munc18-1^A297H^-emGFP or Munc18-1^T304H^-emGFP for 48 h, and then homogenized in 10 mM Tris/Cl, pH 7.5, 150 mM NaCl, 0.5 mM EDTA, 0.5 % NP-40 and protease inhibitors (1/200, Cocktail Set III, EDTA-Free – Calbiochem, Millipore). Tagged proteins were immunoprecipitated using GFP-Trap beads (GFP-Trap®_MA, Chromotek) according to the manufacturer’s instructions, and bound proteins were eluted and analyzed by SDS-PAGE and western blotting with antibodies against VAMP2 (Synaptic Systems, 104202, 1:1000), Munc18-1 (Becton Dickson, 610336, 1:5000) and syntaxin-1A (Sigma-Aldrich, S0664, 1:2000). Blots were visualized and quantified using the Li-Cor Odyssey system (Li-Cor Biosciences). This experiment was performed three times.

#### Immunofluorescence labelling and confocal imaging

DKO-PC12 cells were co-transfected with syntaxin-1A-mEos2 and either Munc18-1^WT^-emGFP, Munc18-1^A297H^-emGFP or Munc18-1^T304H^-emGFP, and then seeded on glass-bottom culture dishes (Cellvis). Cells were washed in Buffer A (145 mM NaCl, 5 mM KCl, 1.2 mM Na_2_HPO_4_, 10 mM D-glucose, 20 mM HEPES, pH 7.4) prior to fixation in 4 % paraformaldehyde in 1x DPBS for 30 min. The cells were then quenched for 10 min (quenching buffer: 50 mM NH_4_Cl in 1x DPBS), permeabilized for 5 min (permeabilization buffer: 0.1 TX-100 in 1x DPBS) and blocked for 10 min (blocking buffer: 0.2 % bovine serum albumin, 0.2 % fish skin gelatin in 1x DPBS) prior to immunolabelling. The cells were incubated with a mouse anti-syntaxin primary antibody (Abcam, ab3265, 1/100) for 60 min at room temperature and then with a goat anti-mouse Alexa 647 secondary antibody (Thermo Fisher Scientific, 1/500) for 45 min in the presence of DAPI (Sigma). The cells were washed 3 times with 1x DPBS for 5 min each wash in between each labelling step. The dishes were washed briefly with Milli Q H_2_0, before mounted with ProLong^TM^ Gold antifade reagent (ThermoFisher) and sealed with a coverslip. For imaging, a laser scanning inverted LSM510 (Zeiss) confocal microscope was used, with a Plan-Apochromat 63x/1.40 NA oil objective.

#### TIRF microscopy

For live-cell TIRF microscopy, transfected cells were visualized using an iLas^2^ ring-TIRF laser illumination system (Roper Scientific) mounted on a Nikon Ti-E inverted microscope, with a 100x/1.49 NA oil-immersion TIRF objective (CFI Apochromat, Nikon) and an Evolve 512 Delta EMCCD camera (Photometrics). Image acquisition was performed using MetaMorph (version 7.10.1.161, Molecular Devices).

#### Secretory vesicle tracking

DKO-PC12 cells were co-transfected with NPY-emGFP and pCMV, Munc18-1^WT^-mCherry, Munc18-1^A297H^-mCherry or Munc18-1^T304H^-mCherry, and then seeded on glass-bottom culture dishes (MatTek). The cells were bathed in Buffer A (145 mM NaCl, 5 mM KCl, 1.2 mM Na_2_HPO_4_, 10 mM D-glucose, 20 mM HEPES, pH 7.4). Time-lapse movies were captured by TIRF microscopy at 20 fps at 37 °C. 1200 frames were acquired before and 3600 frames after the addition of 2 mM BaCl_2_ with the aforementioned frame rate. Particle tracking of NPY-emGFP-labeled SVs was performed on the extracted TIRF images as previously described (Nair et al., 2013). To image vesicles on the plasma membrane, we used a 491 nm laser (Cobolt Calypso, Cobolt Lasers) with an average power of 0.55 mW, or 35-40 % of the maximum power available at the specimen plane, and a dual-band filter set (LF488/561-A-000, Semrock) composed of a dual-band exciter (FF01-482/563-25), dual-band dichroic mirror (Di01-R488/561-25×36) and dual-band emitter (FF01-523/610-25).

#### SV release assay

DKO-PC12 cells were double transfected with VAMP2-pHluorin and Munc18-1^WT^-mCherry, or Munc18-1^A297H^-mCherry, or Munc18-1^T304H^-mCherry. Cells were bathed in Buffer A (145 mM NaCl, 5 mM KCl, 1.2 mM Na_2_HPO_4_, 10 mM D-glucose, and 20 mM HEPES, pH 7.4) and imaged with 2 mM BaCl_2_ stimulation for 10 min (100 ms x 6000 frames). Alternatively, cells were bathed in low K^+^ buffer (0.5 mM MgCl_2_, 2.2 mM CaCl_2_, 5.6 mM KCl, 145 mM NaCl, 5.6 mM D-glucose, 0.5 mM ascorbic acid, 0.1% BSA and 15 mM HEPES, pH 7.4, 290– 310 mOsm), then stimulated and imaged with high K^+^ buffer (56 mM KCl, 0.5 mM ascorbic acid, 0.1% BSA, 15 mM HEPES, 5.6 mM D-glucose, 95 mM NaCl, 0.5 mM MgCl_2_, and 2.2 mM CaCl_2_; at pH 7.4, 290–310 mOsm) for 10 min (100 ms x 6000 frames). SV release was determined by the fluorescence intensity (FI) fold change of VAMP2-pHluorin 200 s after stimulation.

#### Photoactivated localization microscopy (PALM)

Transfected DKO-PC12 cells expressing Munc18-1^WT^-mEos2, Munc18-1^A297H^-mEos2 or Munc18-1^T304H^-mEos2 were incubated in Buffer A alone or stimulated with 2 mM Ba^2+^ for 2 min. The cells were then fixed with 4 % paraformaldehyde diluted in phosphate buffered saline (PBS) for 45 min and washed in PBS. mEos2 molecules were simultaneously photo-converted with a 405 nm laser and excited using a 561 nm laser, as described above. For each dataset, 20,000 frames were acquired at a rate of 33 Hz, at which point the majority of mEos2-tagged molecules had been photoconverted. The acquired 20,000 frames were then processed using Zen 2012 SP5 (Black) (64 bit) Release version 14.0.0.0 (Carl Zeiss Microscopy GmbH) to reconstruct the super-resolution map of each dataset.

#### Single particle tracking PALM (sptPALM)

Time-lapse TIRF movies were captured at 50 Hz (16,000 frames) at 37 °C for both control and stimulated cells. BaCl_2_ (2 mM) was added immediately prior to initiating image acquisition to stimulate the cells. For PALM, a 405 nm laser (Stradus 405, Vortran Laser Technology) was used to photoactivate FP in the cells expressing mEos2-tagged constructs and a 561 nm laser (Cobolt Jive, Cobolt Lasers) was simultaneously used for excitation of the resulting photo converted molecules. To isolate the mEos2 signal from auto fluorescence and background signals, we used a double-beam splitter (LF488/561-A-000, Semrock) and a double-band emitter (FF01-523/610-25, Semrock). To spatially distinguish and temporally separate the stochastically activated molecules during acquisition, the power of the 405 nm laser was adjusted to 1-5 % of the maximum laser power (100 mW) and the 561 nm laser power was adjusted to 75 % of the maximum laser power (150 mW).

### QUANTIFICATION AND STATISTICAL ANALYSIS

#### Secretory vesicle tracking

Briefly, the combination of wavelet segmentation ^64^ and simulated annealing was used to detect and track each molecule. PALM-Tracer ^65^, a custom-written program that runs as a plug-in in the Metamorph software (Molecular Devices), was used to localize single SVs and quantify the SV dynamics. The diffusion coefficient ^60^ distribution was sorted into two groups. The first group, composed of molecules with a D value lower than 0.0312 μm^2^/s, was referred to as “immobile”. The D_threshold_ = 0.0312 μm^2^/s was calculated as previously described ^66^. The second group was defined as the mobile fraction and composed of molecules with D values above 0.0312 μm^2^/s. The average mobile fraction was plotted to assess the changes which occurred in the mobility of vesicles during secretagogue stimulation.

#### Spatiotemporal Cluster Analysis

PALMtracer files were converted using custom MATLAB and python scripts to tryxt and ascii file formats for further processing. Single-particle trajectories of Munc18-1 were drift corrected using SharpViSu ^67^ for spatiotemporal clustering analysis with BOOSH, an experimental variant of the NASTIC suite of software ^44^. Analysis was performed using default parameters for our 50 Hz imaging acquisitions (trajectory steplength >8 steps, ε = 35 nm and MinPts = 3, no MSD filtering, clusters with radius > 150nm excluded). Quantitative comparisons of BOOSH metrics generated from non-stimulated and stimulated conditions were performed using NASTIC Wrangler ^44^.

#### PALM data analysis

The acquired 20,000 frames were then processed using Zen 2012 SP5 (Black) (64 bit) release version 14.0.0.0 (Carl Zeiss Microscopy GmbH) to reconstruct the super-resolution map of each dataset. During processing, every detected fluorescent event was fitted into a two-dimensional Gaussian function to determine the x and y coordinates of its centroid position. After drift correction using inbuilt settings in the software, single-molecule coordinates were compiled to produce the x-y coordinate table for each dataset, from which the corresponding super-resolution image was reconstructed. Cluster analysis was performed using a series of scripts written in Python 2.7. DBSCAN identifies clusters in large datasets of points by a propagative method based on two parameters: r, the search radius and ε, the minimum number of neighbors. It links the points complying with parameters r and ε to propagate clusters. Each point needs to have at least ε number of neighbors within a circle of radius r centered on that molecule in order to be associated with a cluster. The unassigned points are classified as noise. For each cell, three regions of interest (ROIs), each containing between 10000 and 20000 detections, were manually selected. DBSCAN was performed using Python SciKit Learn (sklearn.cluster DBSCAN) to assign points to clusters, with the Convex Hull algorithm (scipy.spatial ConvexHull) used to determine cluster boundaries and areas. Each ROI was analyzed over a range of ε (10-100, stepsize 10) and minPts (2-10, stepsize 1) values. For each ε/minPts analysis a number of DBSCAN-based metrics were returned: total area of the ROI; total detections per ROI; total nanoclusters observed per ROI; nanoclusters per 1000 detections; nanoclusters per μm^2^; percentage of detections in nanoclusters; average number of detections per nanocluster; average nanocluster area; combined area of nanoclusters; percentage of the ROI area in nanocluster; average nanocluster radius (assuming circularity); average density of detections in nanoclusters and average density of all detections in the ROI detections/nm^2^. For each cell the average value of each metric across 3 ROIs represented one statistical n. Comparative analysis was performed using ε = 30 and min Pts = 5, as these values have been shown in our laboratory to return DBSCAN metrics best matching synthetic data generated at the same scale and cluster size/density as experimental data (data not shown). For each experimental condition, outliers for a given metric were removed using median filtering, and the data visualized using Python MatPlotLib. The statistical significance of differences between experimental conditions was determined using independent two-tailed t-tests (scipy. stats t-test_ind), assuming unequal variance.

#### sptPALM analysis

Single-particle tracking data analysis was performed as previously published ^25^. PALM-Tracer ^65^, a custom-written program that runs as a plug-in in Metamorph software (Molecular Devices), was used to localize the molecules and quantify the dynamics of the proteins from 16,000 frame TIRF movies. Each cell was analyzed independently and the distribution of the diffusion coefficients was computed from at least 1000 trajectories. Median values for the MSD of each analyzed cell were calculated from at least 1000 trajectories. The average value of the median was plotted against time. The mobile fraction from the diffusion coefficient distribution for each individual cell was plotted to illustrate cell-to-cell variability. The diffusion coefficient ^60^ distribution was sorted into two groups. The first group, composed of molecules with a *D* value <0.0312 µm^2^/s, was referred to as immobile. The *D*_threshold_ = 0.0312 µm^2^/s was calculated as previously described ^66^. The second group was defined as the mobile fraction and composed of molecules with *D* values higher than 0.0312 µm^2^/s. The color-coding for the superresolved images was performed using ImageJ. The color-coding of the trajectory maps are arbitrary units of 16 colors. The color gradient represents the time of the detection of the tracks in the video. Blue trajectories indicate detection early in the video, whereas pink indicates the late trajectories. In the average-intensity maps, each pixel indicates an individual molecule. The area with the highest density is represented in black, whereas white represents regions with no detection. The color-coding of the diffusion maps ranges from 15,000 units to 50,000, corresponding to diffusion coefficients from 10^−5^ to 10 µm^2^/s.

#### Hidden Markov modeling

We used the variational Bayes SPT (vbSPT) tool ^68^ to analyze Munc18-1^WT^-mEos2, Munc18-1^A297H^-mEos2 and Munc18-1^304H^-mEos2 trajectories and infer motion parameters using a four-state model for statistical comparisons. Cells with more than 1000 trajectories were analyzed as a separate dataset, whereas cells with fewer than 1000 trajectories were pooled together such that the pooled dataset had a minimum of 1000 trajectories.

### DATA AND SOFTWARE AVAILABILITY

The data will be made available upon request.

For the PALM data analysis the acquired 20,000 frames were processed using Zen 2012 SP5 (Black) (64 bit) release version 14.0.0.0 (Carl Zeiss Microscopy GmbH) to reconstruct the super-resolution map of each dataset. DBSCAN was performed using Python SciKit Learn (sklearn.cluster DBSCAN) to assign points to clusters, with the Convex Hull algorithm (scipy.spatial ConvexHull) used to determine cluster boundaries and areas. The data was visualized using Python MatPlotLib. The statistical significance of differences between experimental conditions was determined using independent two-tailed t-tests (scipy. stats t-test_ind), assuming unequal variance. PALM-Tracer ^65^, a custom-written program that runs in Metamorph software (Molecular Devices), was used for the SV tracking and for the sptPALM data analysis. For the Hidden Markov modeling we used the variational Bayes SPT (vbSPT) tool ^68^.

